# Slow delivery immunization enhances HIV neutralizing antibody and germinal center responses via modulation of immunodominance

**DOI:** 10.1101/432666

**Authors:** Kimberly M. Cirelli, Diane G. Carnathan, Bartek Nogal, Oscar L. Rodriguez, Jacob T. Martin, Amit A. Upadhyay, Chiamaka A. Enemuo, Etse H. Gebru, Yury Choe, Federico Viviano, Catherine Nakao, Matthias Pauthner, Samantha Reiss, Christopher A. Cottrell, Raiza Bastidas, William Gibson, Amber N. Wolabaugh, Mariane B. Melo, Benjamin Cosette, Venkatesh Kuman, Nirav Patel, Talar Tokatlian, Sergey Menis, Daniel W. Kulp, Dennis R. Burton, Ben Murrell, Steven E. Bosinger, William R. Schief, Andrew B. Ward, Corey T. Watson, Guido Silvestri, Darrell J. Irvine, Shane Crotty

## Abstract

The observation that humans can produce broadly neutralizing antibodies (bnAbs) against HIV-1 has generated enthusiasm about the potential for a bnAb vaccine against HIV-1. Conventional immunization strategies will likely be insufficient for the development of a bnAb HIV vaccine and vaccines to other difficult pathogens, due to the significant immunological hurdles posed, including B cell immunodominance and germinal center (GC) quantity and quality. Using longitudinal lymph node fine needle aspirates, we found that two independent methods of slow delivery immunization of rhesus macaques (RM) resulted in larger GCs, more robust and sustained GC Tfh cell responses, and GC B cells with improved Env-binding, which correlated with the development of ~20 to 30-fold higher titers of tier 2 HIV-1 nAbs. Using a new RM genomic immunoglobulin loci reference sequence, we identified differential IgV gene usage between slow delivery immunized and conventional bolus immunized animals. The most immunodominant IgV gene used by conventionally immunized animals was associated with many GC B cell lineages. Ab mapping of those GC B cell specificities demonstrated targeting of an immunodominant non-neutralizing trimer base epitope, while that response was muted in slow delivery immunized animals. Thus, alternative immunization strategies appear to enhance nAb development by altering GCs and modulating immunodominance of non-neutralizing epitopes.

## INTRODUCTION

A majority of licensed vaccines provide protection through the induction of protective antibodies (Plotkin, 2010). The isolation of HIV-1 broadly neutralizing antibodies (bnAbs) from numerous HIV-infected individuals, combined with passive transfer studies demonstrating that HIV-1 bnAbs can protect non-human primates (NHPs) from SHIV infections, supports the feasibility of an antibody-based HIV vaccine (Burton and Hangartner, 2016; Nishimura and Martin, 2017). Elicitation of neutralizing antibodies (nAbs) against clinically relevant HIV-1 strains (i.e., tier 2 and tier 3 strains) by immunization, however, has been very difficult (Montefiori et al., 2018). Much of that challenge centers on structural features of HIV-1 envelope (Env); those structural features have complex and incompletely understood immunological implications. HIV-1 Env consists of gp120 and gp41 components that form a trimeric spike that is the only surface protein on HIV-1 virions and thus is the only possible target for nAbs (Burton and Hangartner, 2016). Immunization of humans with monomeric HIV-1 gp120 has repeatedly failed to elicit tier 2 nAbs in candidate HIV-1 vaccine clinical trials (Haynes et al., 2012; Mascola et al., 1996; Rerks-Ngarm et al., 2009). The reasons for that are not intuitively obvious, as nAb epitopes are present on monomeric gp120. Immunodominance of non-neutralizing epitopes is one possible explanation (Havenar-Daughton et al., 2017). After decades of effort, key protein design developments have been made in recent years to accomplish expression of soluble native-like HIV-1 Env trimers (Julien et al., 2013; Kulp et al., 2017; Lyumkis et al., 2013; Sanders et al., 2013). In contrast to gp120, immunization with native-like Env trimers elicited substantial strain-specific tier 2 nAbs in rabbits and guinea pigs (Feng et al., 2016; Sanders et al., 2015). However, immunization with native-like Env trimers failed to elicit tier 2 nAbs in mice (Hu et al., 2015), and Env trimer immunizations of NHPs have only been sporadically successful (Havenar-Daughton et al., 2016a; Pauthner et al., 2017; Sanders et al., 2015; Zhou et al., 2017). For some immunization regimes in NHPs, tier 2 nAbs have been reliably elicited within 10 weeks of Env trimer immunization (Pauthner et al., 2017), which compares favorably with the speed of tier 2 nAb development in HIV infected individuals (Richman et al., 2003; Wei et al., 2003). Thus, while nAb epitopes are clearly presented on native-like Env trimers, the immunological parameters controlling the development of nAbs to the Env trimer remain to be elucidated. These immunological parameters are also likely important for nAbs to other pathogens.

Germinal centers (GCs) are essential for HIV-1 nAb development, as HIV-1 nAb development requires antibody (Ab) somatic hypermutation (SHM) (Klein et al., 2013; West et al., 2014). GCs are sites where B cells compete for antigen and undergo repeated rounds of SHM of their BCRs and selection by GC T follicular helper cells (Tfh) to evolve high affinity Abs (Crotty, 2014; Mesin et al., 2016). B cells with higher affinity to antigen present more peptide:MHC complexes to GC Tfh cells and in turn receive more help (Crotty, 2014; Gitlin et al., 2014; Victora et al., 2010). GC Tfh help signals to GC B cells result in proliferation and further BCR mutation (Gitlin et al., 2015). Additional parameters may also regulate competition in GCs and rates of SHM. Tfh help quality was associated with HIV nAb development in Env trimer immunized rhesus macaque monkeys (RM) (Havenar-Daughton et al., 2016a). In HIV-infected humans, frequencies of highly functional memory Tfh cells in blood were associated with bnAb development (Locci et al., 2013; Moody et al., 2016). HIV-specific circulating Tfh (cTfh) were positively correlated with total HIV-specific Ab development (Baiyegunhi et al., 2018). GC Tfh cells were also positively correlated with nAb development in SIV^+^ RMs and SHIV^+^ RMs (Chowdhury et al., 2015; Petrovas et al., 2012; Yamamoto et al., 2015).

Affinity maturation is only one component of nAb development. B cell responses to protein antigens are polyclonal, targeting epitopes across an antigen. The composition of the antigen-specific B cell repertoire responding to even a single protein can be complex. The responding B cells initially engage in interclonal competition, and then the B cells develop numerous GCs and engage in interclonal and intraclonal competition, resulting in complex outcomes (Kuraoka et al., 2016; Tas et al., 2016). Theoretically, the entire surface of any protein represents a continuum of B cell epitopes. In reality, the Ab response to a protein predominantly targets a limited number of epitopic sites. This phenomenon is well described for influenza HA, and the epitopes are recognized in a hierarchical manner (Angeletti and Yewdell, 2018; Angeletti et al., 2017). Immunodominance is the phenomenon in which B cells that recognize an epitopic site dominate an immune response at the expense of B cells that recognize other sites. Immunodominance can occur due to differences in B cell precursor frequencies and affinities (Abbott et al., 2018; Havenar-Daughton et al., 2017; Jardine et al., 2016) and appears to be a key immunological process limiting the development of broad nAb responses to influenza (Andrews et al., 2018; Angeletti and Yewdell, 2018; Angeletti et al., 2017; Victora and Wilson, 2015). Immunodominance may also be important for the development of nAbs against refractory pathogens, including HIV-1. Evidence of immunodominance impairing HIV nAb development includes the lack of tier 2 nAb responses by humans immunized with Env gp120, the lack of tier 2 nAb responses in RMs immunized with non-native Env trimers, the sporadic nature of tier 2 nAb development in RMs, and the role of immunodominance in the response of rare or low affinity HIV CD4-binding-site specific B cells in a mouse model (Abbott et al., 2018; Havenar-Daughton et al., 2017; 2018).

Much of the focus in HIV vaccine development is on the choice of antigen and adjuvant, but an orthogonal parameter is the kinetics of the availability of the antigen. Slow, or sustained, delivery immunization is a conceptually attractive vaccine strategy because it more closely mimics a natural self-limiting acute infection (Cirelli and Crotty, 2017). While the adjuvanticity of alum has been believed to be in part due to a ‘depot’ effect of sustained antigen availability, many antigens rapidly elute from alum in vivo (Hogenesch, 2002; Shi et al., 2001; Weissburg et al., 1995) and several studies reported that the depot attribute of alum did not affect Ab responses (Hogenesch, 2012; Hutchison et al., 2012; Noe et al., 2010), suggesting that alum adjuvanticity does not primarily function via a slow antigen release mechanism. In contrast, two-week slow release immunization using nonmechanical osmotic minipumps and a soluble adjuvant resulted in enhanced GC B and Tfh cell responses in mouse models (Hu et al., 2015; Tam et al., 2016). Two-week dose escalation immunization using a soluble adjuvant resulted in similar outcomes and enhanced deposition of immune complexes onto follicular dendritic cells (FDC) (Tam et al., 2016).

Current understanding of the relative importance of different aspects of B and T cell biology in the development of HIV nAbs has been limited by the fact the wildtype mice do not develop tier 2 HIV nAbs in response to Env trimer immunization. While NHPs are important animal models for HIV vaccine design because of their close evolutionary relationship to humans, it has been very difficult to study the early response to Env in NHPs, and humans, due to the inaccessible nature of lymph nodes (LN) and the low frequencies of Env-specific B cells in response to a primary immunization. In a first NHP slow release vaccine study, six RMs were immunized with soluble native-like Env trimers in a soluble ISCOMs-class saponin adjuvant delivered via nonmechanical osmotic pumps to test the concept of slow release immunization (Pauthner et al., 2017). The minipump immunized animals responded with the most robust tier 2 nAb responses of any of the groups of animals immunized. Tier 2 nAb responses were developed by wk10 in all minipump immunized RMs. The rapidity and peak magnitude of the tier 2 nAb response suggested that improved affinity maturation, altered B cell lineage recruitment, enhanced Env-specific Tfh cell responses, or other factors may be responsible for the improved nAb response. Antigen-specific B cell and Tfh cells were not examined.

We considered that comparison of the primary B and T cell responses in the draining LNs of osmotic minipump immunized RMs and conventionally immunized RMs may provide insights into the immunological causes of the difficulty in eliciting B cell responses capable of neutralizing tier 2 HIV strains, which may also be applicable to other difficult-to-neutralize pathogens. Here we have examined the early B and T cell response to HIV Env trimers in RMs using new tools and comparative immunology between conventional and slow release vaccine concepts to gain insights into the development of nAbs.

## RESULTS

### Env-specific GC responses are more robust upon slow release immunization

Three groups of RMs were immunized with soluble native-like Env trimer BG505 Olio6-CD4ko (Kulp et al., 2017) protein in a soluble ISCOMs-class saponin adjuvant. Three delivery strategies were tested: conventional bolus immunization via subcutaneous (SubQ) needle injection (n = 9), two-week SubQ nonmechanical osmotic minipumps (n = 4) and four-week SubQ nonmechanical osmotic minipumps (n = 4) (**Fig 1A**). All immunizations were given bilaterally in left and right thighs. To determine the kinetics of the GC response to primary immunization, longitudinal LN fine needle aspirates (FNA) were employed. LN FNAs were used to sample the draining inguinal LNs weekly in both the left and right leg. Previous work demonstrated that LN FNAs well represented the cellular composition of the whole LN and were well tolerated (Havenar-Daughton et al., 2016a). This study is the first longitudinal (i.e., same individuals sampled) weekly kinetic analysis of a GC response in any species.

**Figure 1.**
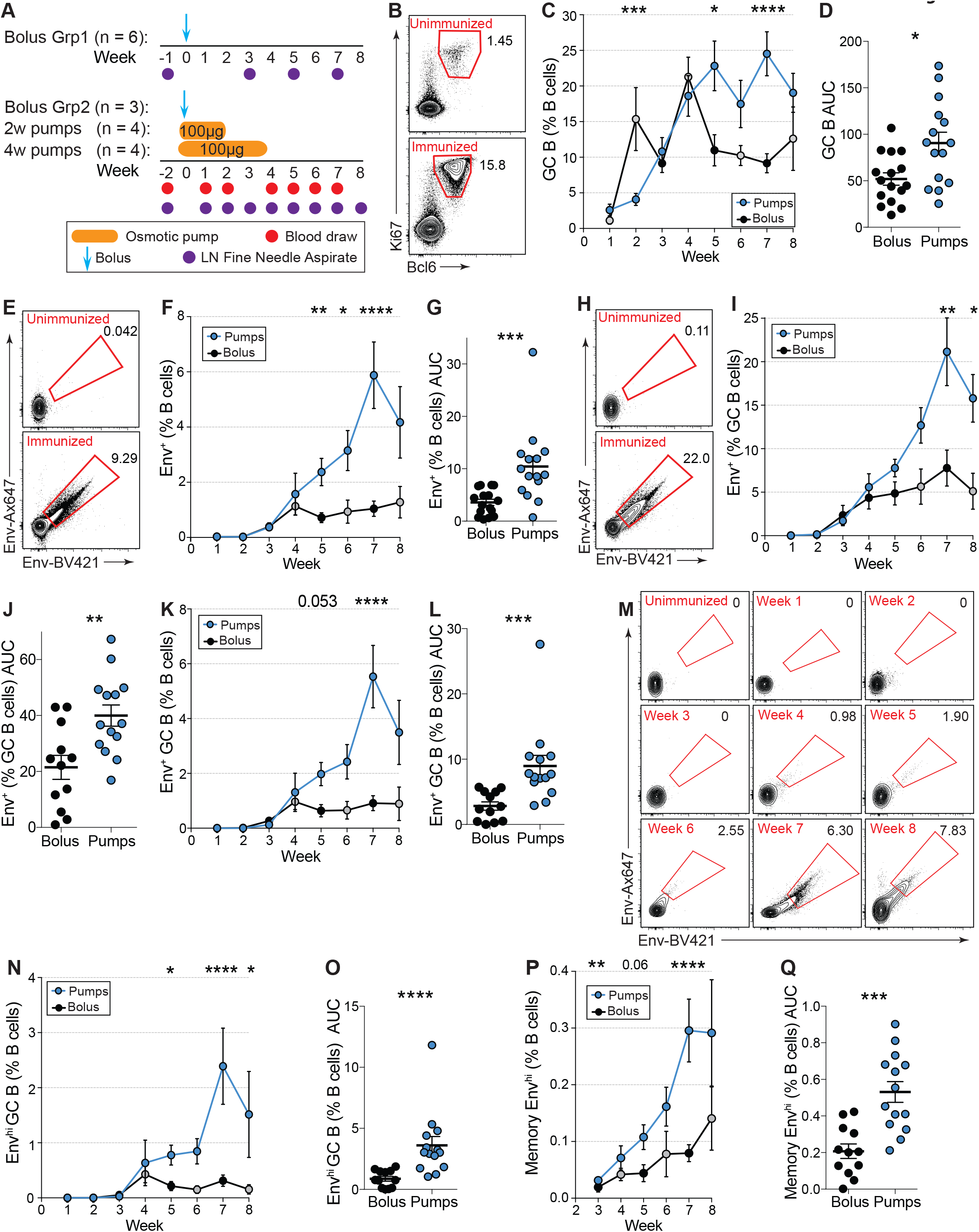
Sustained delivery immunization enhances germinal center (GC) B cell responses. (A) Immunization and sampling schedule of first immunization. Bolus Grp2, 2w pumps, and 4w pump groups were immunized and sampled at the same time. Bolus Grp1 were immunized and sampled at a later time. Bolus Grps 1 and 2 data have been pooled. (B) Representative flow cytometry gate of GC B cells, gated on viable CD2O^+^ B cells pre- and post-immunization. See **Fig S1** for full gating strategy. (C) GC B cell frequencies over time. Black circles are time points when bolus groups have been pooled (n = 9), grey circles (n =6). 2 week pump and 4 week osmotic pump groups have been pooled in all analyses. GC B cells were quantified as Bcl6^+^ Ki67^+^ at weeks 1–6, and 8. At week 7, GC B cells were defined as CD38^−^ CD71^+^ (see **Fig S1** for gating). (D) Cumulative GC B cell responses to immunization within individual LNs at weeks 1, 3–7 [AUC]. (E) Representative flow cytometry gate of Env-specific B cells pre- and post-immunization. Env-specific cells are gated as Env_AX647_^+^ EnvBV42i^+^ (IgM^+^ IgG^+^)^−^ CD2O^+^ CD3^−^ cells. (F) Env-specific B cell frequencies over time. (G) Cumulative Env-specific B cell responses within individual LNs at weeks 1, 3–7 [AUC]. (H) Representative flow cytometry gating of Env-specific GC B cells pre- and post-immunization. Cells are gated as Env_AX647_^+^ Env_BV421_^+^ of Bcl6^+^ Ki67^+^ (IgM^+^ IgG^+^)^−^ CD20^+^ CD3^−^ or Env_AX647_^+^ Env_BV421_^+^ of CD38^−^ CD71^+^ (IgM^+^ IgG^+^)^−^ CD2O^+^ CD3^−^. (I) Quantification of Env-specific GC B cells, quantified as percentage of total GC B cells, over time. (J) Cumulative Env-specific GC B cell responses within individual LNs at weeks 1, 3–7 [AUC]. (K) Quantification of Env-specific GC B cells, quantified as percentage of total B cells, over time. (L) Cumulative Env-specific GC B cell responses within individual LNs at weeks 1, 3–7 [AUC]. (M) Flow cytometry gate of high-affinity Env-specific GC B cells over one immunization within an individual LN. (N) Frequencies of high-affinity Env-specific GC B cell, quantified as percentage of total B cells, over time. (O) Cumulative high-affinity Env-specific GC B cell responses within individual LNs at weeks 1, 3–7 [AUC]. (P) Quantification of high-affinity memory B cells over time. Memory B cells were calculated as non-GC (Bcl6^−^ Ki67^−^ or CD38^+^ CD71^−^) high-affinity Env-specific B cells. (Q) Cumulative high-affinity memory B cell responses within individual LNs at weeks i, 3–7 [AUC]. Mean ± SEM are graphed. Statistical significance tested using unpaired, two-tailed Mann-Whitney U tests. *p≤0.05, **p≤0.01. ***p≤0.001, ****p≤0.0001

GCs developed slower than expected after conventional bolus immunization, based on comparison to mouse data of LN GC kinetics after protein immunization, with almost no GC B cells (Bcl6^+^Ki 67^+^ or CD38^−^ CD71^+^ of CD20^+^CD3^−^) detectable at day 7 (d7) postimmunization (**Fig 1B-C, S1A - B**). Substantially greater GC B cell frequencies were present at d14 (d7 v d14, p= 0.0015). No differences were observed in GC kinetics between the two osmotic pump groups, so all data from those animals have been pooled in subsequent analyses (n = 8 animals; n = 16 LN FNAs per time point. **Fig S1C-D**). Total GC B cells in the draining LNs peaked at week 7 (w7) in pump-immunized animals after a single immunization, substantially later than after bolus immunization (**Fig 1C**). Pump-immunized animals had significantly more GC B cells throughout the first immunization (p = 0.017 [Area under the curve (AUC)]. **Fig 1D, S1D**).

Given that RMs are not kept in a sterile environment, interpretation of GC B cell kinetics, in the absence of antigen-specific probes, is confounded by uncertainty regarding the antigenic targets of the GCs. In previous studies, total GC B cell responses were measured, but antigen-specificity was not determined (Havenar-Daughton et al., 2016a; Pauthner et al., 2017). Detection of antigen-specific GC B cells is a particular challenge, as GC B cells express less BCR than non-GC B cells (**Fig S1E**). Here, using BG505 Olio6 Env trimers conjugated to two fluorochromes as two separate and complementary Env trimer probes (Env_A647_ and Env_BV421_), we measured the kinetics and magnitude of the Env trimer-specific B and GC B cell response (Env_A647_^+^ Env_BV421_^+^ Bcl6^+^Ki67^+^ or CD38^−^ CD71^+^ of CD20^+^CD3^−^) (**Fig 1E-O, S1F-N**). This method is specific, with little experimental ‘noise’, as naive B cells and GC B cells from unimmunized animals did not bind these probes (**Fig 1E, H, M, S1G, S1K, Table S1–2**). Despite observing considerable GCs at weeks 2–3, Env-specific GC B cells with detectable affinity to the probes were rare at weeks 2–3 (**Fig 1I-K, S1G**). Antigen-specific B and GC B cells in draining LNs of bolus immunized animals were consistently detectable at w4. Env-specific GC B cell frequencies were relatively stable between weeks 4–8 in bolus immunized animals, indicating active GC responses for at least two months after a single protein immunization (**Fig 1I, 1K**).

In minipump-immunized animals, frequencies of Env-specific GC B cells increased over time (p = 0.0064 compared to bolus as Env^+^ % of GC B cells over time [AUC], and p = 0.0001 compared to bolus as Env^+^ GC B cell % of total B cells over time [AUC]. **Fig 1H-L**). Enhanced GC B cell binding of Env by pump immunized animals was not due to an increase in BCR expression (**Fig S1H**). MFI of B cell binding to Env can be used as a surrogate metric of binding affinity (**Fig 1M, S1I, S1J**). High affinity Env-specific GC B cells became much more abundant in minipump immunized animals over time (p < 0.0001 compared to bolus over time [AUC], **Fig 1N-O, S1I-N**), suggesting that osmotic minipump vaccine administration resulted in more affinity maturation compared to conventional bolus immunization.

Env trimer-specific memory B cells (Bcl6^−^ Ki67^−^ or CD38^+^ CD71^−^ Env_Ax647_^+/hi^ Env_BV421_^+/hi^ CD20^+^ cells) developed in draining LNs in response to conventional or slow release immunization (**Fig 1P-Q, S1O-P**). Minipump immunized animals developed significantly higher frequencies of high affinity Env trimer-specific memory B cells (**Fig 1P-Q**). Overall, these GC and memory B cell data demonstrate that slow immunization delivery resulted in more robust GCs and indicated substantially greater affinity maturation to Env after a single immunization than occurred upon conventional bolus immunization.

### Slow release osmotic minipumps enhance Env-specific GC Tfh cell responses

While total GC Tfh cell (CXCR5^+^ PD1^hi^ of CD4^+^ CD8^−^) frequencies were significantly increased in pump animals at several time points during the first immunization, overall GC Tfh cells did not differ between groups (**Fig 2A-C, S2A**). The specificity of GC Tfh at these time points could not be measured due to limited cell number recoveries and experimental prioritization of the B cell assays.

**Figure 2.**
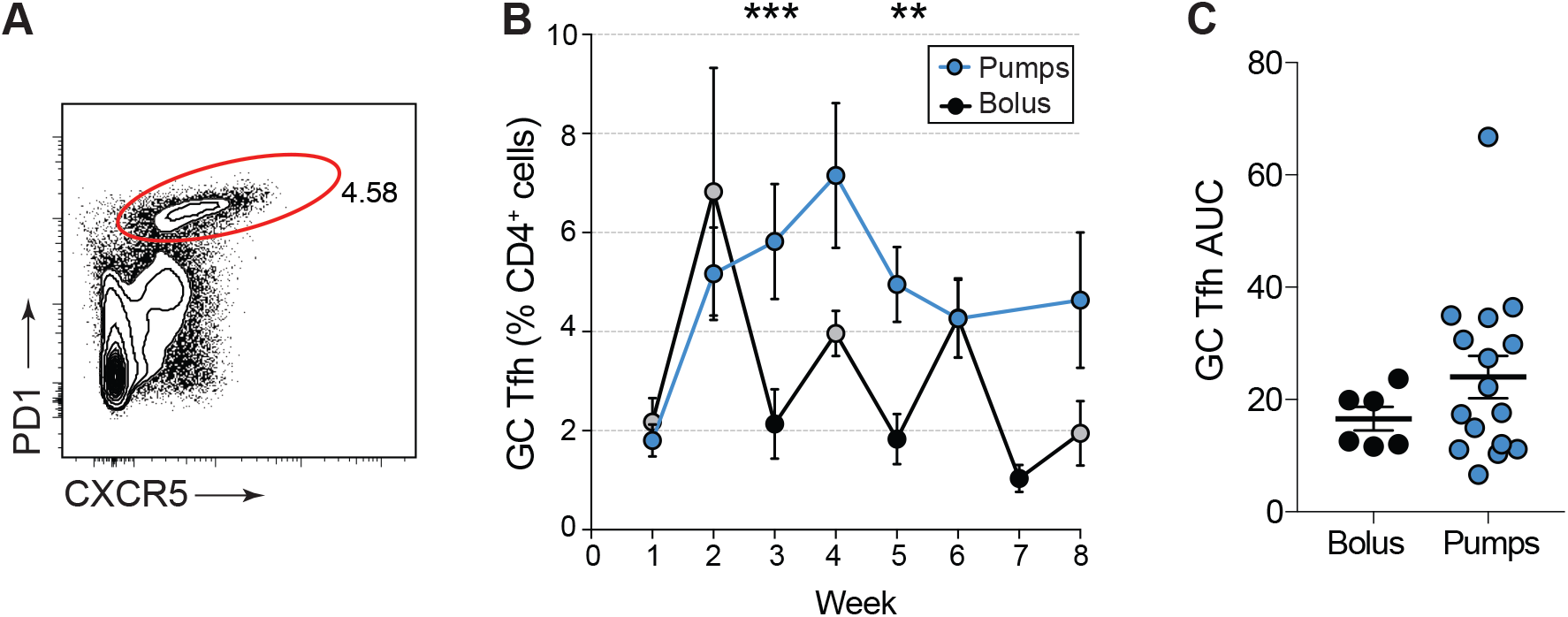
Sustained delivery immunization enhances GC Tfh responses. (A) Representative flow cytometry gate of GC Tfh, gated on CD4^+^ T cells. See **Fig S2** for full gating strategy. (B) Quantification of GC Tfh cells over time. (C) Cumulative GC Tfh cell response to Env immunization between at weeks 1, 3–6 [AUC]. Mean ± SEM are graphed. Statistical significance tested using unpaired, two-tailed Mann-Whitney U test. **p≤0.01. ***p≤0.001

Based on previous immunization regimens (Pauthner et al., 2017), we administered a 2^nd^ Env trimer immunization at w8 (**Fig 3A**). For minipump immunized animals, the immunization was split evenly between osmotic pumps and a bolus administered at the end of pump delivery to simulate an escalating dose immunization. We hypothesized that a bolus immunization at the end of the slow release delivery may enhance plasma cell differentiation and Ab titers. The total dose of Env trimer was matched between groups (100μg, **Fig 3A**). Draining LN GC responses observed after the 2^nd^ immunization were relatively flat (**Fig 3B**), perhaps because the GC responses were already well above baseline immediately prior to the 2^nd^ immunization (**Fig 1C, 3B**), though other explanations are also possible (see Discussion). Minipump immunized animals had significantly larger GC B responses at w14 (**Fig 3B**). Env-specific B and GC B cell frequencies of bolus immunized animals increased after the 2^nd^ immunization (**Fig 3C-F**). High affinity Env-specific GC B cell recall responses were largely comparable between slow release immunized animals and bolus immunized animals (**Fig 3F**). Overall, kinetics of the secondary GC responses differed from those in the primary GC responses.

**Figure 3.**
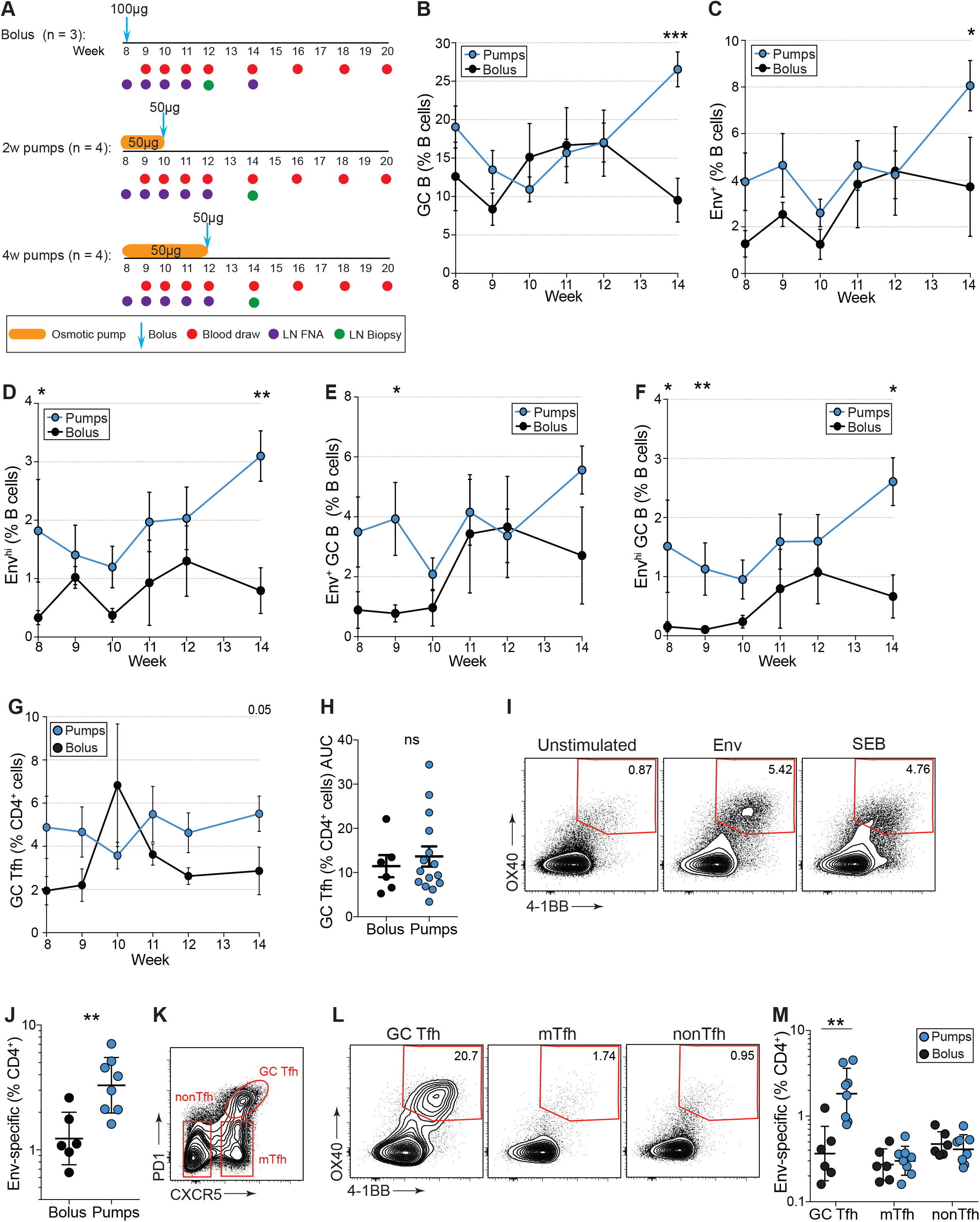
Germinal center responses following 2^nd^ Env trimer immunization. (A) Immunization and sampling schedule of 2^nd^ immunization. All groups were immunized and sampled contemporaneously. (B) Frequencies of total GC B cells over time, gated as per **Fig 1B**. (C) Env-specific B cell frequencies over time, gated as per **Fig 1E** and **S1F**. (D) Frequencies of high-affinity Env-specific B cells over time, gated as per **Fig S1I**. (E) Quantification of Env-specific GC B cells over time, as gated per **Fig 1H** and **S1G**. (F) Quantification of high-affinity Env-specific GC B cells over time, as gated per **Fig 1M** and **S1K**. (G) GC Tfh frequencies after second immunization, gated as per **Fig 2A**. (H) Cumulative GC Tfh cell responses in response to the Env booster immunization within individual LNs between weeks 9 and 12 [AUC]. (I) Representative flow cytometry plots of Env-specific CD4 T cells, gated on viable CD4^+^ T cells. LN cells were left unstimulated or stimulated with a pool of overlapping peptides spanning Olio6-CD4ko (Env). SEB is shown as a positive control. Frequencies are background-subtracted (J) Quantification of Env-specific CD4^+^ T cells at week 12 (bolus) or week 14 (pumps). (K) Representative flow cytometry gating of GC Tfh, mantle (m)Tfh and nonTfh subsets. (L) Flow cytometry plots of AIM_OB_ assay, gated on GC Tfh, mTfh or nonTfh cells. (M) Quantification of Env-specific CD4^+^ T cells by subset. Mean ± SEM are graphed. Statistical significance tested using unpaired, two-tailed Mann-Whitney U test. *p≤0.05, **p≤0.01, ***p≤0.001

Total GC Tfh cell frequencies were similar in response to the 2^nd^ Env trimer immunization (**Fig 3G-H**). To identify Env-specific GC Tfh cells, we performed cytokine-agnostic activation induced marker (AIM) flow cytometry assays with biopsied LN cells (Dan et al., 2016; Havenar-Daughton et al., 2016b; Reiss et al., 2017). Higher frequencies of Env-specific CD4^+^ T cells were present in slow release immunized animals compared to bolus immunized animals (**Fig 3I-J, S2B**). The Env-specific CD4^+^ T cell response enhancement was selective to Env-specific GC Tfh cells (**Fig 3K-M**). In conclusion, slow release immunization delivery elicited an immune response that generated substantially more Env-specific GC Tfh cells, commensurate with the development of significantly higher frequencies of high affinity Env-specific GC B cells.

### Slow release osmotic pump delivery enhances humoral responses

Antibody responses to the different immunization approaches were examined, in light of the differential GC responses detected. A single bolus immunization failed to elicit detectable BG505 Env trimer-specific serum IgG titers, (**Fig 4A, S3A-B**). In contrast, a single slow release minipump immunization elicited modest but significant Env trimer-specific plasma IgG titers (w7, p = 0.048). The Olio6-CD4ko Env trimer design included a His tag; the tag elicited a strong anti-His Ab response after a single minipump immunization (**Fig 4B, S3C-D**), while bolus immunized animals made anti-His IgG responses after the booster immunization. A fraction of the Env-specific B and GC B cells likely recognized the His epitope. The 2^nd^ Olio6-CD4ko Env trimer immunization induced anamnestic Env IgG responses in both the conventional bolus immunized animals and the slow release minipump immunized animals, with minipump outperforming conventional bolus immunization (**Fig 4A**). BG505 Env-specific IgG titers increased in response to the 2^nd^ minipump immunization prior to the end-of-regimen bolus injection, demonstrating that slow delivery immunization alone was sufficient for substantial anamnestic plasma cell development (w7 vs w10, p = 0.008). Env-binding IgG titers between conventional bolus groups and between osmotic minipump groups were similar to the previous study after each immunization (**Fig S3E**).

**Figure 4.**
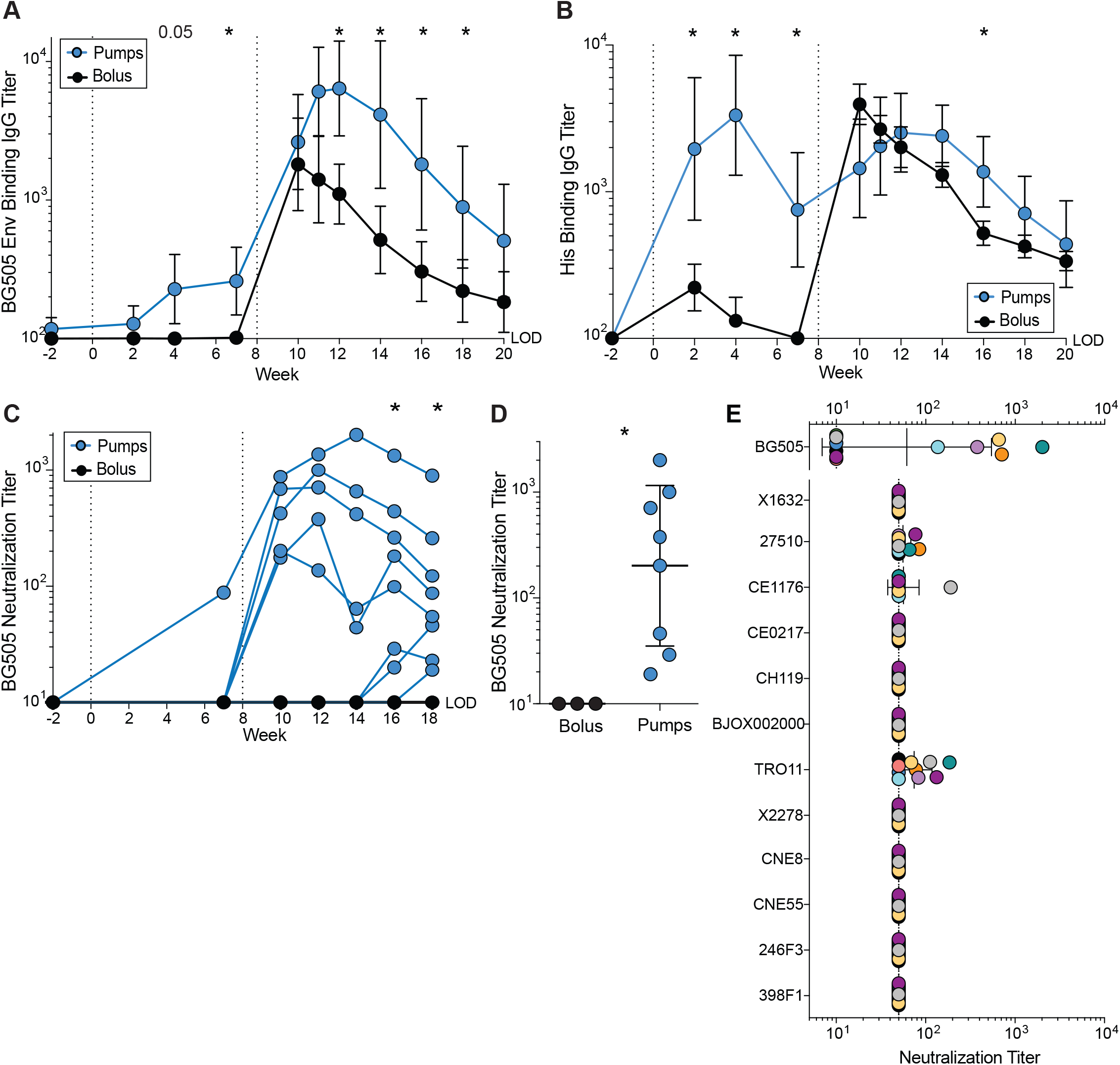
Sustained delivery immunization induces higher nAb titers than conventional immunization. (A) Env (BG505) trimer binding IgG titers over time. (B) Anti-his IgG binding titers over time. (C) BG505 N332 nAb titers over time. (D) Peak BG505 N332 nAb titers after two immunizations. (E) Neutralization breadth on a 12 virus panel, representing global antigenic diversity, at week 10 (bolus), 12 (2w pumps) and 14 (4w pumps). All data represent geometric mean titers ± geometric SD. Statistical significance tested using unpaired, two-tailed Mann-Whitney U test. *p≤0.05

To assess the development of autologous tier 2 nAb titers over time, sera were tested for neutralization of BG505 N332 pseudovirus using TZM-bl neutralization assays (**Fig 4C-D, S3F-H**). By w10, 5/8 osmotic minipump immunized animals developed nAbs in contrast to 0/3 bolus animals (1:99 vs < 1:10 geometric mean titer [GMT]). All minipump immunized animals developed nAbs by w18 (**Fig 4C**). Peak BG505 neutralization titers of minipump animals were ~20-fold higher than bolus animals (1:202 vs 1:10 GMT, p = 0.01). 6/8 minipump immunized animals demonstrated partial neutralization breadth, neutralizing one to three heterologous HIV-1 isolates (**Fig 4E, S3J**). No heterologous tier 2 nAbs were detected in animals that received a conventional immunization regimen. In sum, slow release immunization enhanced the magnitude and quality of the Ab response to Env immunization, which was associated with the enhanced Env-specific GC B cell and GC Tfh cell responses.

### Slow delivery immunization alters the antigen-specific B cell repertoire

Because of the higher frequencies of high-affinity B cells and nAb titers observed in the minipump immunized animals, we hypothesized that slow release immunization delivery may affect several aspects of B cell responses. Firstly, slow antigen delivery may activate (direct effect) or recruit (via T cell help) more diverse B cell lineages. Inclusion of more independent clonal lineages of B cells would increase the likelihood that B cells with rarer and/or lower-affinity BCRs capable of developing into nAbs will be expanded. Secondly, slow delivery immunization may result in the generation of higher numbers of memory B cells capable of re-circulating and reseeding new GCs among multiple LNs upon booster immunization. Finally, slow release immunization may drive higher frequencies of SHM. A major technical challenge for testing these hypotheses in NHPs was the lack of a complete genomic sequence of the RM immunoglobulin (Ig) gene loci. A complete germline Ig gene reference is required for proper B cell lineage assignment and identification of authentic SHM. While a RM genome sequence was available (Gibbs et al., 2007), the Ig genes were largely unmapped because Ig genes reside within highly complex genomic regions that are characterized by high levels of repetitive sequence architecture and inter-individual haplotype variation (Watson et al., 2017). Genomic characterization and annotation of Ig genes has proven challenging because of this complexity (Watson and Breden, 2012). Most next generation sequencing techniques use short read technologies (~150bp), which can be insufficient for resolving large (>15kb) repetitive segmental duplications (Alkan et al., 2011). Therefore, we sequenced the genome of a RM using Pacific Biosciences (PacBio) long-read sequencing technology to 60-fold coverage. Overall, reads obtained had a median length of 16.6kb and a maximum length of 69.4kb. Genome assembly was conducted using FALCON/FALCON-Unzip (Chin et al., 2016), resulting in a total of 1,633 primary contigs with a median length of 8.4mb (2.83gb total bases).

Contigs containing the IGH, IGL and IGK loci were identified, and V, D and J genes were annotated via a combination of bioinformatics and manual curation (**Fig 5A**). 66 IGHV, 41 IGHD, 6 IGHJ, 68 IGKV, 5 IGKJ, 65 IGLV, and 7 IGLJ genes were identified by focusing on gene segment annotations with open reading frames (ORFs; **Fig 5A**). Notably, the long reads generated from this experiment allowed for the characterization of regions that were unresolved in previous assemblies, including the current RM reference genome (rheMac8), facilitating descriptions of novel gene loci (**Fig 5B**). On top of identification of novel gene loci, it was possible to identify heterozygous allelic variants at loci identified in primary contigs by using a combination of raw PacBio read data and alternate contigs from FALCON-Unzip, facilitated by the long reads (**Fig 5C-E**). Together, we determined that 37/66 IGHV, 31/68 IGKV, and 12/65 IGLV genes were heterozygous, amounting to a germline database of 103, 99, and 77 V alleles for each respective locus (**Fig 5E, Table S3**). Sequencing BCR RNA of mature B cells from the same animal and close relatives supported the presence of these annotated ORF sequences (data not shown). A significant fraction of alleles identified in the PacBio assemblies were not represented by sequences in either the IMGT database or NCBI repositories, highlighting the utility of this approach for improving upon existing genomic databases (**Fig 5F**). In contrast to previous work suggesting differences in Ig gene sub-family composition between human and RM (Vigdorovich et al., 2016), we found gene family sizes to be comparable between the two species (**Fig 5G**).

**Figure 5.**
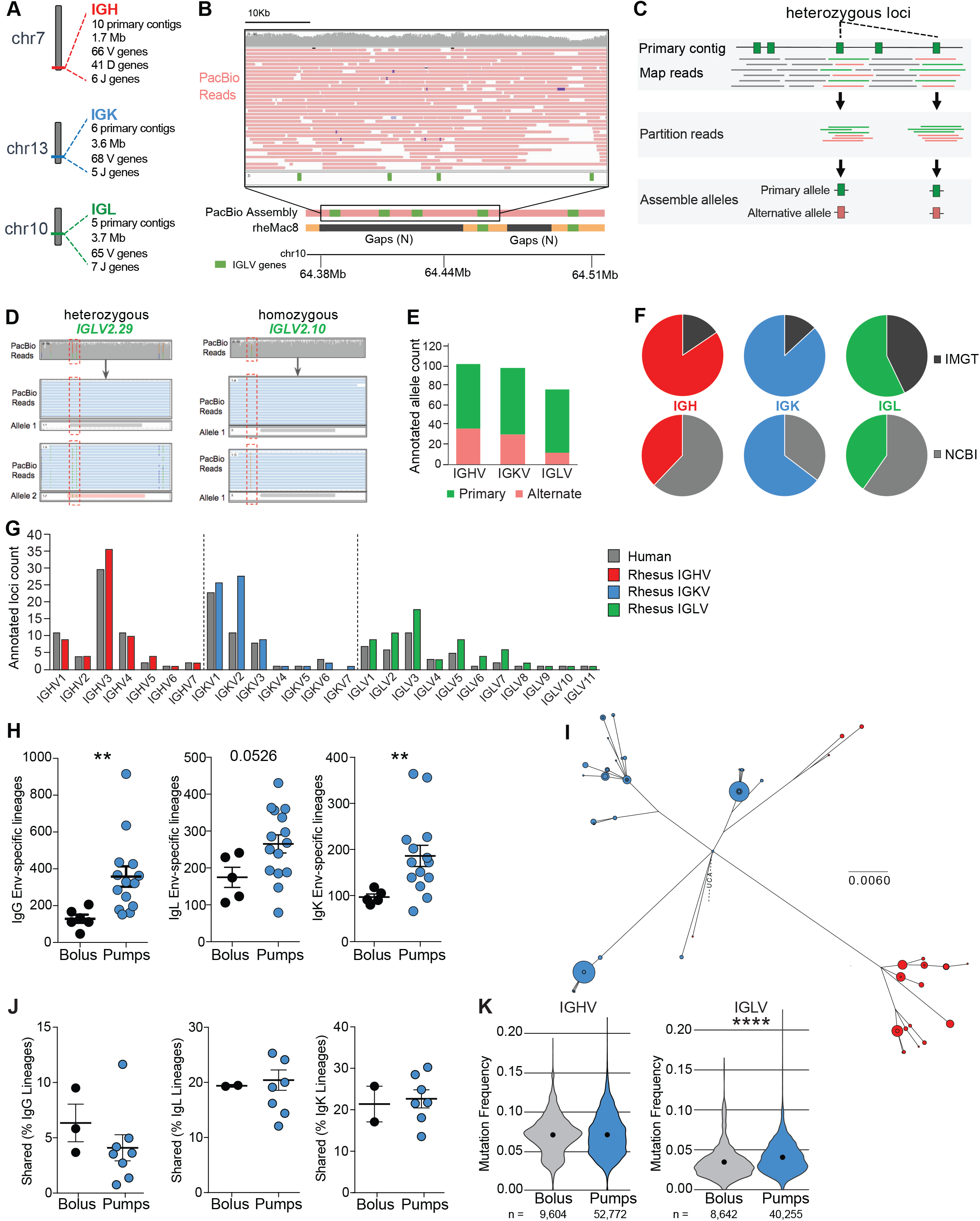
Immunoglobulin germline annotations using long-read genomic DNA sequencing. (A) Locus and high-level assembly summaries for the three primary Ig loci in RM. (B) An example region in the IGL locus, in which PacBio primary contig assemblies resolved existing gaps in the current RM reference genome (rheMac8). Contig sequences spanning these gaps resulted in an increase in the number of known mapped IGLV loci. The inset provides an example of PacBio reads representing the current assembly that span the largest of these gaps, revealing high and consistent read coverage, with multiple reads spanning three of the novel genes displayed. (C) Overview of V gene allelic variant discovery process. PacBio reads overlapping ORF annotations on primary contigs were assessed for the presence of SNPs. At ORF loci determined to be heterozygous, SNPs were used to partition reads for local allele-specific assemblies. (D) SNPs within and near genes (red boxes) were used to partition PacBio reads to each respective haplotype, allowing for the identification of heterozygous and homozygous gene segments. Following this approach, an additional alternative *IGLV2.29* allele was found by mapping PacBio reads to the primary contigs (left); in contrast, reads overlapping *IGLV2.10* provide evidence for the presence of identical alleles on both homologous chromosomes. (E) Counts of annotations from primary contigs and alternate alleles identified from PacBio reads using the method described in (C) and (D). (F) The proportions of IGHV, IGKV, and IGLV genes/alleles annotated from PacBio assembly data that are present in IMGT and NCBI repositories. (G) Counts of IGHV, IGKV, and IGLV gene loci (partitioned by subfamily) annotated from the current PacBio primary contig assemblies, compared to the equivalent counts of known gene loci in human. (H) Quantification of total number of IgG, IgL and IgK lineages of Env-specific B cells. Each data point is an individual LN. Env-specific B cells were sorted from bolus immunized animals at week 12 and pump immunized animals at week 14. (I) Phylogenetic analysis of an Ag-specific lineage found in both LNs in a single animal. Blue, left LN; Red, right LN. Size of dot represents number of reads with that sequence. (J) Percentage of lineages shared between R and L LN within a given animal. (K) Violin plots of mutation frequencies in (B) IGHV or (C) IGLV. Black dot is mean. Mean ± SEM are graphed. *p≤ 0.05, **p≤0.01, ***p≤0.001

To assess how slow delivery immunization affected the repertoire of the Env-specific B cell response, we isolated and sequenced BCRs from Env-specific B cells in the draining LNs of animals immunized with conventional or slow release modalities (**Fig S4A**). The majority of the sequenced Env-specific B cells were GC B cells (77%), providing a window into this difficult-to-study cell type. Utilizing the new RM Ig reference sequence, we assigned each unique BCR sequence to the V and J genes with most similarity, performed lineage analysis, and determined SHMs. More Env-specific B cells were isolated from minipump immunized animals compared to bolus animals (303,644 vs. 52,302 cells [total]; 20,242 vs. 8,717 cells [mean], p = 0.029) (**Table S1**), consistent with the higher frequencies of Env-specific B cells identified by flow cytometry (**Fig 1, 3**). Much greater numbers of unique Env-specific BCR sequences were isolated from slow delivery immunized animals than bolus animals, both for heavy chain (52,772 vs 9,604 *IgG*) and light chains (40,255 vs 8,642 *IgL;* 39,131 vs 6,358 *IgK*). Furthermore, significantly more unique *IgG* and *IgK* B cell lineages were identified in LNs of minipump immunized animals compared to conventional bolus immunized animals (**Fig 5H, S4B**). While most BCR lineages were found in only one LN (**Fig S5C**), 0.7 – 30.2% were found in both R and L LNs (**Fig 5I-J**). SHM rates in minipump and bolus BCR lineages were largely similar (**Fig 5K, S4D-E**). Thus, substantially more Env-specific GC B cell lineages were sustained in animals receiving a slow release immunization, while SHM rates were comparable.

### Slow delivery immunization resulted in greater diversity of antibodies and recognized epitopes

Given that slow release immunization resulted in more Env-specific B cell lineage diversity, we sought to determine if differential IgV gene usage occurred, which may suggest differences in the epitopes targeted on the Env trimer. Strikingly, bolus animals utilized IGLV3.15 and IGHV3.76 significantly more frequently than pump animals (q = 0.00003 and q = 0.03, respectively) (**Fig 6A-B**). 21.9% and 13.5% of Env-specific B cells from LNs of bolus animals utilized IGLV3.15 and IGHV3.76, respectively. 3.3% and 4.1% of Env-specific B cells from LNs from minipump animals utilized IGLV3.15 and IGHV3.76, respectively. Using IMGT for similar analysis, a difference in IGLV3.15 (aka IGLV3-10) was identified between groups (**Fig S4F**). No difference in IGHV gene use was identified due to the low number of V genes available in IMGT. Analyzing the data with a broader Ig database incorporating both genomic and RNA sequencing data (Corcoran et al., 2016; Ramesh et al., 2017; Sundling et al., 2012), use of IGLV3.15 (aka IGLV3-5) was again significantly higher among bolus immunized animals (q = 0.0003) (**Fig S4G**). E nv-specific B cells that used IGLV3.15 were phylogenetically diverse and could be found in both draining LNs within a single animal (**Fig 6C, S5A**).

The differential use of IGLV3.15 suggested that the Env-specific B cells elicited by bolus immunization targeted epitopes distinct from the Env-specific B cells elicited by slow release immunization. Taken together with the lack of HIV-1 nAbs in the conventional bolus immunized animals, we hypothesized that B cells that utilized IGLV3.15 recognized the base of the trimer. This region is normally hidden on full length Env expressed on virions. In contrast, the base is the largest proteinaceous region exposed on soluble Env trimer due to the unusually dense glycans covering most of the remainder of the surface of HIV-1 Env (Stewart-Jones et al., 2016)(**Fig 6D**). The base is a major non-neutralizing Ab target in mice and macaques immunized with soluble Env trimer, and base-specific B cells are proposed to be immunodominant to nAb-epitope-specific B cells (Havenar-Daughton et al., 2017; Hu et al., 2015; Kulp et al., 2017). To test this hypothesis, we sequenced 196 Env-specific single B cells from the draining LNs of two bolus immunized animals at w7 to obtain paired BCR sequences utilizing IGLV3.15. We selected an IGLV3.15 utilizing clone and synthesized the corresponding monoclonal Ab (mAb), termed BDA1. The IGLV3.15 utilizing mAb BDA1 bound BG505 Env trimer, but not monomeric BG505 gp120 or His peptide. (**Figure 6E-F, S5B**). BDA1 binding to Env trimer was selectively blocked by 19R, a high affinity Env base-binding mAb, demonstrating that BDA1 recognizes the Env trimer base (**Figure 6G**). EM analysis of a BDA1 Fab complex with BG505 Env trimer confirmed binding of BDA1 to the trimer base (**Figure 6H**). We next sought to determine how BDA1 (w7) was related to the Env-specific B cells isolated from the same LN after booster immunization with Env trimer (w12). Alignment and phylogenetic analysis of the BDA1 lineage consisted of BDA1, three related w12 sequences, and the inferred germline sequence, with few mutations between the BDA1 heavy chain and the related w12 IgG GC B cell sequences (**Fig S5C-D**). The BDA1 IGLV3.15 light chain displayed more diversity between w7 and w12, indicating recall GC responses of IGV3.15^+^ cells and ongoing SHM (**Fig S5E-F**).

**Figure 6.**
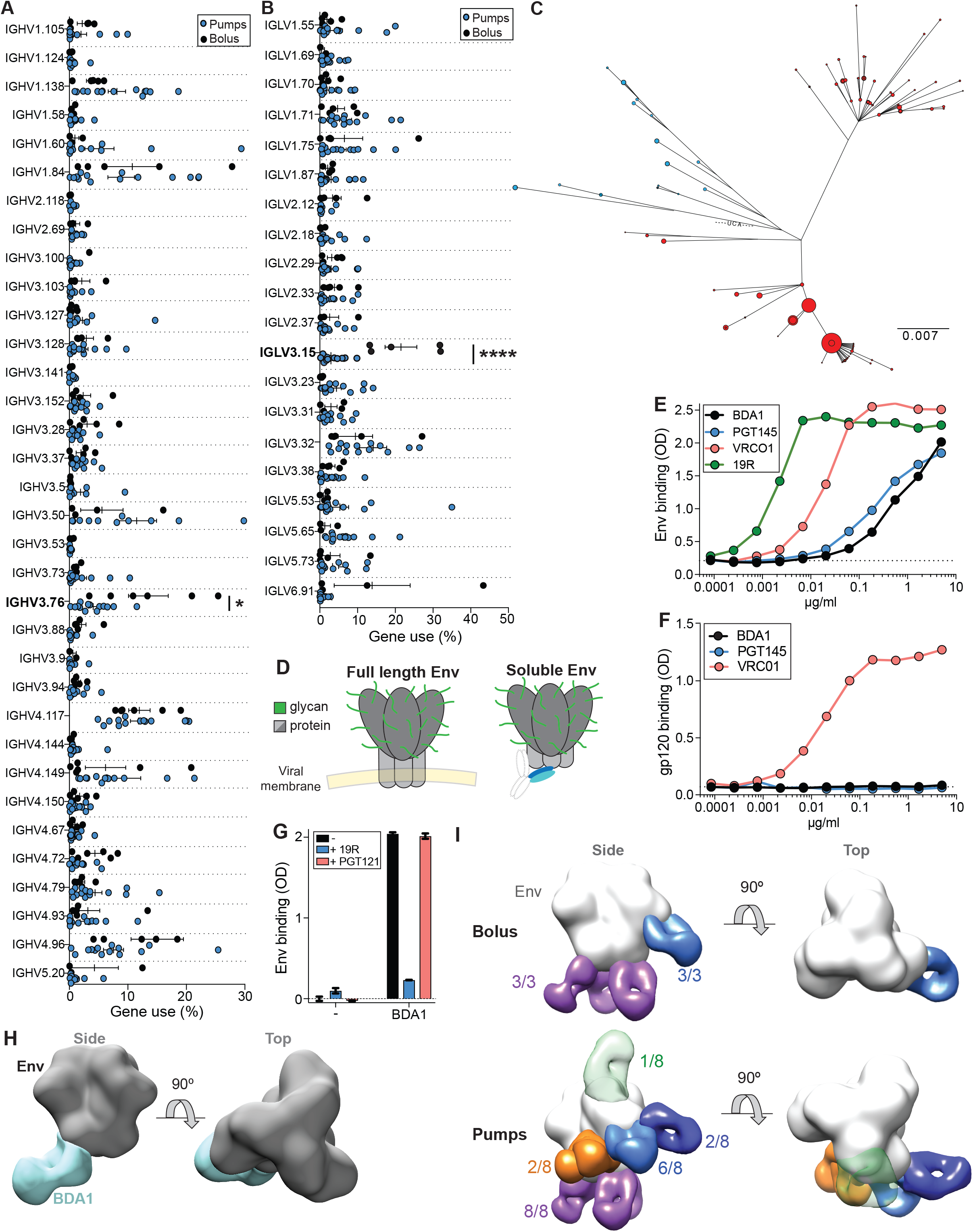
Sustained immunization shifts immunodominance. Percentage of (A) IGHV or (B) IGLV use by antigen-specific B cells within a lymph node. Each data point is single LN. Mean ± SEM are graphed. *q<0.05, ****q<0.0001, FDR = 5%. (C) Phylogenetic tree of a single IGLV3.15-utilizing lineage. Blue, left LN; Red, right LN. Size of dot represents number of reads with that sequence. (D) The base of the Env trimer is not exposed on the surface of the virion. Soluble trimer used in immunizations allows access of the base to B cells. Glycans on the surface of the trimer restrict access of B cells to proteinaceous surface. (E) Binding curves of BDA1 and bnAbs to BG505 Env trimer. (F) Binding curves of BDA1 and bnAbs to gp120 monomer. (G) Cross-competition ELISA assay. BDA1, base-directed antibody utilizing IGLV3.15 isolated from a bolus immunized animal at w7. 19R is a base-binding antibody isolated from an immunized RM. PGT121 is a bnAb targeting the N332 epitope towards the top of the trimer. 19R fab, PGT121 fab and BDA1 whole antibody were used in this assay. Data shown are representative of two experiments, each performed in duplicate. (H) 3D EM reconstruction of BDA1 Fab (blue) in complex with BG505 SOSIPv5.2 Env trimer. (I) Composite 3D reconstruction of Env trimer bound to Fabs isolated from sera of all animals after two immunizations, as determined by polyclonal EM analysis. Numbers of individual animals with Fab that binds region are listed. Base (purple), N355 (light blue), C3/V5 (dark blue), fusion peptide (orange), apex (green). Apex specific fab is depicted as transparent because fabs with this specificity were rare, but present. 3D EM reconstructions for plasma antibodies from individual animals can be seen in **Figure S7**.

We utilized polyclonal EM serological analysis as an independent approach to assess the Ab responses to Env trimer between the two immunization strategies (Bianchi et al., 2018). This new technique allows for simultaneous visualization of diverse Abs targeting distinct epitopes, directly from polyclonal serum processed into Fabs. Ab responses in conventional bolus immunized animals targeted two sites on Env: the trimer base (3/3 animals), and the N335 region (3/3 animals). (**Fig 6I, S6**). In contrast, the polyclonal Ab responses in minipump immunized animals were substantially more diverse. In addition to the base and N335 regions, three potential nAb epitopes, the fusion peptide, V1/V3, and C3/V5 regions (Klasse et al., 2018; Kong et al., 2016) were targeted by pump immunized animals (**Fig 6I, S6**). Base directed Ab responses were present in minipump immunized animals, as expected. In sum, BCR sequencing, mAb characterization, and polyclonal EM analyses demonstrated that slow release immunization resulted in a substantial shift in the GC B cell and Ab response towards Env epitopes that are both distinct from, and more diverse than, the Env epitopes predominantly targeted in response to conventional bolus immunization. Notably, the shifted response is towards nAb epitopes, which are likely immunorecessive epitopes in comparison to the epitopes of the Env trimer base, indicating that slow release immunization causes a substantial modulation of immunodominance or change in the immunodominance hierarchy. Together with the Env-specific GC B cell kinetics and the enhanced Env-specific GC Tfh cell responses, these data provide a logical and plausible immunological explanation for the dramatic difference in HIV-1 nAb titers between the groups.

### Escalating dose immunization enhances germinal center and nAb responses

Dose escalation is an immunization strategy to achieve extended antigen exposure that is an approach distinct from osmotic minipumps (Tam et al., 2016). Escalating dose (ED) immunization has the added advantage of mimicking the antigen dose kinetics of an acute infection. Therefore, an NHP ED study was performed with Env trimer as an independent assessment of the immunological implications of extended (two week) antigen delivery in a vaccine setting. The control group was given conventional bolus immunizations at w0, w10, and w24, totaling 100μg, 100μg, and 300μg of Olio6 native-like Env trimer protein, respectively, mixed with an ISCOMs-class adjuvant, as per the osmotic minipump study (**Fig 7A**). ED immunizations were administered as 7 injections over 2 weeks (**Fig 7A**), with a total antigen dose equivalent to that of the conventional bolus immunization group. Significantly higher frequencies of GC B cells in draining LNs were observed at w5 in the ED group compared to the conventional bolus immunization group (**Fig 7B-C, S7A**). ED immunization resulted in significantly more Env-specific B and GC B cell after the 1^st^ immunization (p = 0.0002 [AUC]) (**Fig 7D-E, S7B-F, Table S4**). ED immunization also elicited improved affinity maturation, as indicated by the enhanced development of Env^hi^ GC B cells compared to conventional immunization after the 1^st^ immunization (**Fig S7G-J, Table S5**). Additionally, ED immunization resulted in significantly more Env-specific memory B cells compared to conventional immunization after the 1^st^ immunization (**Fig S7K-N**). Total GC B cell frequencies, and Env-specific GC and memory B cell frequencies also increased upon the 2^nd^ and 3^rd^ ED immunizations, though not above the peak frequencies observed in response to the 1^st^ ED regimen (**Fig 7B-E, S7A-N**). Analysis of CD4^+^ T cells in the draining LNs by LN FNA revealed that ED resulted in significantly higher total GC Tfh and Env-specific GC Tfh after the 1^st^ immunization (**Fig 7F-H, S7O-P**). ED immunized animals showed a higher ratio of Env^+^ GC B cells: Env-specific GC Tfh cells, suggesting that ED immunization results in greater antigen-specific help to B cells than conventional immunization (**Fig 7I**). The magnitude of the improved primary Env-specific GC B cell response, the increased GC Tfh cell response, and the enhanced Env^hi^ GC B cell response upon ED immunization were comparable to those observed after minipump immunization.

**Figure 7.**
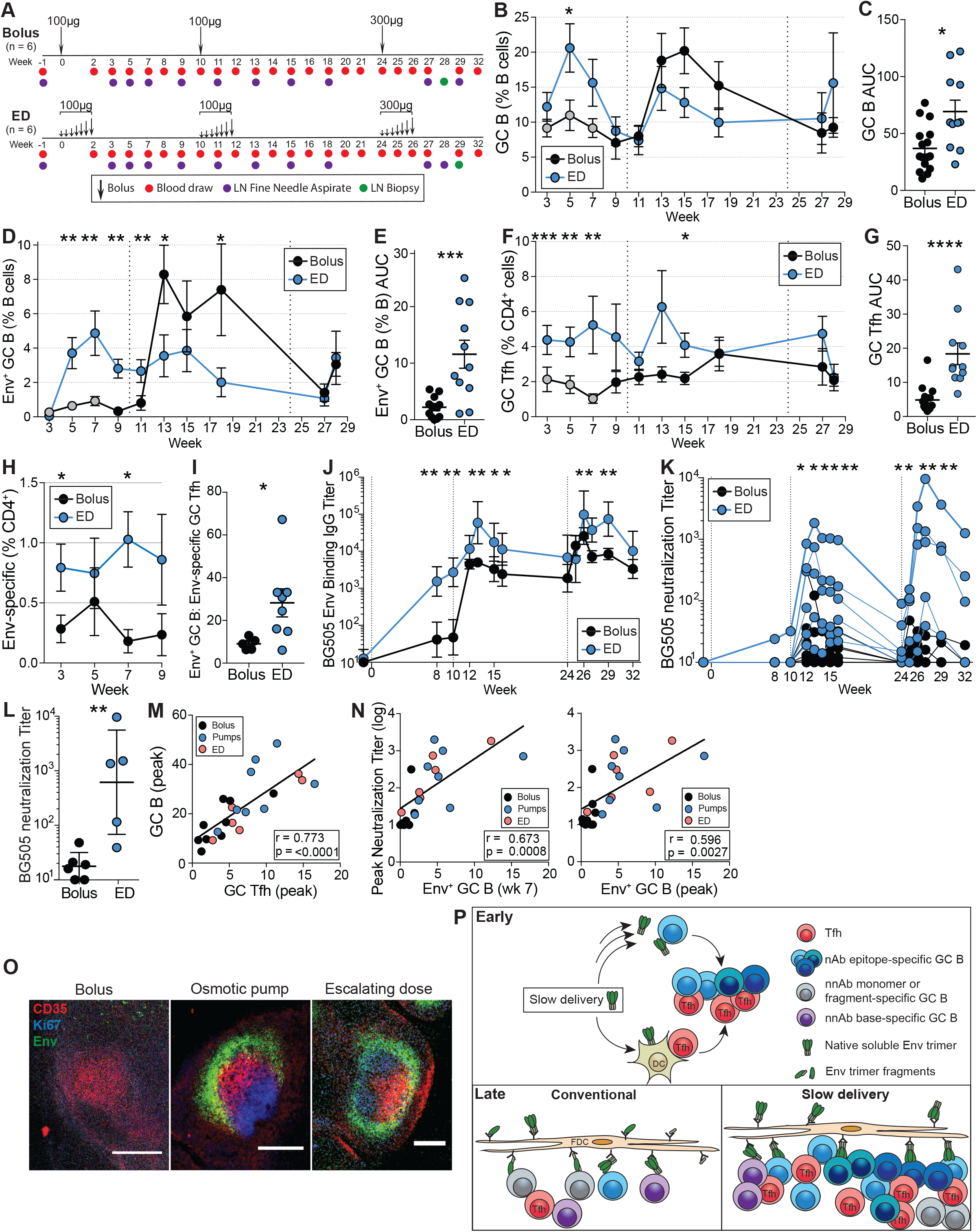
Dose escalating immunization strategy results in higher nAb titers. (A) Immunization and sampling schedule. Groups were immunized and sampled at the same time. (B) Quantification of total GC B frequencies over time. Data from bolus grp2 at w3, 5, and 7 (Fig 1) are included in these analyses (grey circles). (C) Cumulative GC B cell responses to the first immunization [AUC]. AUC was calculated between w3–7. (D) Quantification of Env-specific GC B cells frequencies over time. (E) Cumulative Env-specific GC B cell responses to the first immunization [AUC]. (F) Quantification of total GC Tfh cell frequencies over time. (G) Cumulative GC Tfh responses to the first immunization [AUC]. (H) Quantification of Env-specific CD4^+^ responses after 1 immunization. (I) Ratio of Env^+^ GC B cells to Env-specific GC Tfh at w5, calculated as Env^+^ GC B cells (% B cells)/ Env-specific GC Tfh (% CD4^+^). (J) Total BG505 Env trimer binding IgG titers over time. (K) BG505 N332 nAb titers over time. (L) Peak BG505 N332 nAb titers after three immunizations. (M) Correlation between peak GC Tfh and GC B cells frequencies during 1^st^ immunization. Data is from both studies. (N) Correlation between Env^+^ GC B cells (% B cells) and peak neutralization titers. Env^+^ GC B cell values are from w7 or peak frequencies during 1^st^ immunization. Peak neutralization titers are after 2^nd^ immunization. (O) Histology of inguinal LNs from RMs immunized with Env_AX647_ (Alexa647-labelled MD39) and ISCOMs-class adjuvant via conventional bolus (n = 3), 2 week osmotic pump as per Fig 1A (n = 3) or an escalating dose regimen as per Fig 7A (n = 3). LNs were harvested 2 days after the end of the immunization (bolus, d2; pump, d16; ED, d14). Green, Env; red, CD35; blue, Ki67. Scale bars, 250μm. (P) Model of GC response in conventional immunization vs. slow delivery. Slow delivery immunization likely alters early (~d1-d7) activation and differentiation of Tfh cells and activation and recruitment of a diverse set of B cells. Greater GC Tfh help supports a wider repertoire of B cells, which is more likely to contain nAb precursors, later in the response (w3–7). Antigen delivered via conventional bolus immunization can be subject to degradative processes and nonnative forms of antigen can be presented by FDCs late in the response, while pumps protect the antigen prior to release. IC formation is enhanced to slow delivery immunization. All BG505 binding and neutralization data represent geometric mean titers and ± geometric SD. All cell-frequency data represent mean and SEM. *p<0.05, **p<0.01, ***p<0.001, ****p<0.0001

A single ED immunization regimen was sufficient to elicit a BG505 Env-specific IgG response (**Fig 7J**). Anamnestic Env-binding plasma Ab responses were observed after the 2^nd^ and 3^rd^ DE and conventional immunizations (**Fig 7J**). All ED immunized animals developed tier 2 BG505 HIV-1 nAbs after the 2^nd^ immunization, while only 3/6 conventionally immunized animals developed nAbs (**Fig 7K**). Peak tier 2 nAb titers after the 3^rd^ immunization were ~30-fold higher in ED immunized RMs, significantly greater than conventionally immunized RMs (1:615 vs 1:18 GMT, p = 0.009) (**Fig 7L**; nAb breadth in **Fig S7Q**).

Total GC Tfh frequencies correlated with total GC B frequencies during the 1^st^ immunization (r = 0.773, p = <0.0001 [peak of 1^st^ immunization], **Fig 7M**). In a previous study, total GC B frequencies correlated with nAb development (Pauthner et al., 2017). A primary hypothesis of this study was that the magnitude of Env-specific GC B cell responses to the 1^st^ immunization might predict nAb development. Peak Env^+^ GC B frequencies to the 1^st^ immunization correlated with peak nAb titers in response to the 2^nd^ immunization (r = 0.673, p = 0.0008 [w7]; r = 0.596, p = 0.0027 [peak of 1^st^ immunization]. **Fig 7N**), indicating that Env^+^ GC B cell frequencies and Env-specific GC Tfh cell responses can predict subsequent nAb development.

Taken together, the data show that the ED immunization modality generated greater GC and humoral responses than dose-matched conventional immunization, closely recapitulating the immune responses elicited to osmotic minipump immunization, indicating that modulation of GC B cell and Tfh cell responses is a general property of slow delivery immunization strategies, which can result in dramatically different B cell specificities and nAb development.

An ED regimen resulted in enhanced FDC deposition of antigen in mice (Tam et al., 2016). We hypothesized that the enhanced GC responses observed here in Env trimer immunized RMs with both slow delivery immunization modalities were, at least in part, due to increased availability of antigen to GC B cells and GC Tfh cells (Cirelli and Crotty, 2017). Therefore, an antigen tracking study was performed in RMs with labeled Env trimer and ISCOMs-class adjuvant administered via a conventional bolus (n =3), 2w osmotic pump (n=3) or an escalating dose regimen (n=3). Histological analyses of draining LNs revealed extensive Env localized within follicles after pump or ED immunization, while none was detectable in those of bolus immunized animals (**Fig 7O**). Thus, slow delivery immunization leads to enhanced antigen retention within LNs in NHPs.

## DISCUSSION

Understanding the underlying immunological challenges to nAb development against challenging pathogens may be important for understanding why protective immunity to such pathogens is elusive; direct examination of primary immune responses in lymphoid tissue is required to develop such an understanding. Strategies to enhance the humoral and GC responses to immunization are also likely needed for the development of vaccines against some complex pathogens, particularly HIV-1. Using two independent methods, we have demonstrated that slow delivery immunization resulted in enhanced Tier 2 nAb development in NHPs. To examine the immune responses directly in the draining LNs, we employed weekly LN FNAs. From this, we were able to ascertain that conventional bolus immunization elicited a robust GC response, but that slow delivery immunization altered the kinetics and overall magnitude of the GC response. Strikingly, slow antigen delivery modulated the immunodominance of the B cell response to non-neutralizing epitopes on Env trimer. Each of these differences were prominent during the primary immune response and were positively associated with the larger nAb response that subsequently developed in slow delivery immunized animals, suggesting that much of the failure of a bolus immunization to a difficult antigen is intrinsic to early B cell events associated with immunodominance features of multi-epitope complex antigens.

Detailed weekly analysis of the primary immune response to Env trimer in draining LNs of RMs immunized by conventional bolus injection revealed more durable GC responses than expected, with relatively stable Env-specific GC B cell frequencies from 4–8 weeks postimmunization. Perhaps the most dramatic observation in the bolus immunized animals was the degree of immunodominance observed in the GC B cell response to Env trimer. Approximately 25% of Env trimer-specific B cells in bolus immunized animals were IGLV3.15^+^. An antibody utilizing IGLV3.15, BDA1, targeted the nonneutralizing base of the Env trimer. The base is a major site recognized by Abs of recombinant trimer-immunized animals (Havenar-Daughton et al., 2017). The Env trimer base appears to be immunodominant because it is a large exposed protein surface with many potential epitopes and acceptable BCR angles of approach when compared to the other surfaces of Env trimer, which are predominantly shielded by large glycans. Because BDA1 had relatively modest affinity for Env trimer, and many IGLV3.15^+^ GC B cell lineages were observed, we speculate that the IGLV3.15^+^ base-specific B cell response is immunodominant because the precursor B cells are common rather than being of particularly high affinity. More broadly, differences in B cell responses to the same antigen between immunization strategies, the BDA1 data, Env-specific BCR sequence repertoire results, Env trimer binding Ab titers, nAb titers, and the polyclonal Ab EM mapping together demonstrate substantial immunodominance of non-neutralizing B cells that outcompete B cells specific for neutralizing epitopes after a conventional bolus injection.

The most unexpected outcome was that slow antigen delivery altered the repertoire of the responding Env-specific B cells and the range of Ab specificities and nAb specificities. The simplest explanation for this outcome was that slow antigen delivery increases the likelihood that rare and/or lower affinity immunorecessive nAb precursors are recruited into the B cell response, resulting in more diversity in the epitopes targeted among the GC B cells. We reiterate that the antigen dose was equal between the bolus and pump immunized animals, and between the bolus and ED animals, thus total antigen dose is not the driver of these differential outcomes. A small fraction of Env-specific B cells from minipump immunized animals utilized IGLV3.15, but Abs isolated from pump animals still targeted the base, consistent with diverse epitopes accessible on the base allowing BCRs targeting this site utilizing diverse *IGHV* and *IGLV* genes.

While the trimer base was exposed in both contexts, differences in epitope accessibility on the Env trimer may exist between immunization strategies. Immune complexes (ICs), composed of Env trimer and antibodies, are bound by FDCs for presentation to B cells. Binding of an antibody to its cognate epitope, however, can block further access to that epitope by B cells undergoing selection. We speculate that a large fraction of the early antibody response targets the base, later reducing the response against this region. During a slow delivery immunization, early base-specific nnAbs may form ICs with newly available Env trimers, enhancing presentation on FDCs and possibly increasing the likelihood that nAb epitope-specific GC B cells will be selected for survival both due to increased antigen availability in the GC and the orientation of the Env trimer occluding the base (**Fig 7P**).

We had predicted that slow release immunization would reduce the B cell response to protein breakdown products and fragments that occur in vivo, such as the internal face of gp120 and V3, by protecting the antigen in its native form, thus having a greater percentage of intact Env trimer antigen on FDCs at 2–10 weeks postimmunization (**Fig 7P**) (Cirelli and Crotty, 2017; Tam et al., 2016). However, differential responses to intact Env trimer were not predicted. GC B cell responses to gp120 internal face and breakdown products are surely present in each of these groups of animals. These cells likely make up a substantial fraction of the ‘dark antigen’ GC response (Kuraoka et al., 2016), and may have immunodominant specificities, as a majority of GC B cells did not bind intact native Env trimer with measurable affinity (**Fig 1I, S7G**). Adjuvant alone does not induce a GC B cell response (Havenar-Daughton et al., 2016a), consistent with the conclusion that the GC B cells elicited in these immunizations are predominantly specific for Env. While those specificities are of some interest, in this study we focused on the Env trimer-binding B cells, due to the limited cell numbers in each sample.

As noted above, the simplest explanation for altered immunodominance was increased recruitment of rare or lower affinity immunorecessive nAb precursors into the B cell response. It has been reported that naive antigen-specific B cells normally have a narrow window of time of a few days to be recruited into a GC response (Turner et al., 2017). A narrow window of time for B cell recruitment disproportionately affects B cells with rare precursor frequencies. Slow antigen delivery may substantially expand the pool of recruited B cells by extending that window, thereby increasing the breadth of the B cell repertoire sampled by the draining LN. Additionally, Tfh selection of B cells based on affinity (peptide-MHCII complex presentation) may be most stringent prior to the GC response (Schwickert et al., 2011; Yeh et al., 2018); slow antigen delivery may reduce that stringency by substantially broadening the time window for Tfh interactions with Env-specific B cells of differing epitope specificities at the border of the follicle. The diversity of the B cell response would then likely be greater at the end of the immunization, if the diversity was maintained.

The data are also consistent with multiple mechanisms of action potentially altering the nAb outcomes. Several aspects of GC biology were affected by slow delivery. Slow delivery immunization induced higher frequencies and numbers of total and Env-specific GC Tfh cells. Slow delivery regimens resulted in greater retention of antigen within LNs. Greater availability of GC Tfh cell help and antigen to B cells was accompanied by larger and more enduring GC B responses. Slow delivery resulted in a substantially higher number of Env-specific GC B cells. The GC B cells were also more diverse, as defined by unique Env-specific B cell lineages, which may be a consequence of broader initial activation of antigen-specific B cells described above, or a consequence of sustaining larger GCs over time, or both. The biological relevance of those processes is reinforced by the observation of more diverse nAb Env-binding specificities generated in slow delivery immunized animals contrasted with conventional bolus immunization, as defined by polyclonal Ab EM mapping.

NAb and germinal center responses to secondary immunizations in conventional bolus immunized animals were weaker in this study than a previous study (Pauthner et al., 2017); this may be due to a change in adjuvant formulation. Nevertheless, nAbs were robust in response to minipump or ED immunization.

Despite a considerable apparent difference in affinity maturation (Env^hi^ GC B cells and Env^hi^ memory B cells), SHM rates were largely equivalent between groups after two immunizations. The data suggest that differential rates of SHM were not the cause of the improved affinity maturation and improved nAb responses. A study examining SHM in memory B cells after RM immunizations with nonnative Env trimers and a range of adjuvants (which did not elicit tier 2 nAbs) did not observe differences in SHM between groups (Francica et al., 2015). The SHM data are again consistent with a model where the primary cause of the difference in the neutralizing Ab outcomes was the altered immunodominance profile of the B cell response.

While slow delivery immunization can alter the immunodominance of epitopes and enhance the response to immunization, immunogens should be also be optimized to minimize responses against non-neutralizing epitopes. These results are consistent with immunodominance findings in HIV-1 bnAb mouse models (Abbott et al., 2018; Duan et al., 2018). Osmotic pumps have been used in humans for drug delivery and are feasible for early human vaccine trials. However, it is impractical for large-scale vaccination efforts, as it requires a simple surgery. Nevertheless, ED is technically available immediately as a GC enhancing alternative to conventional bolus immunization. Less cumbersome slow delivery immunization technologies are worthy of further development, including degradable encapsulating biomaterials and depot forming adjuvants that make antigen available over time (i.e., not rendered inert in the depot) in ways that sustain GCs (DeMuth et al., 2013; 2014). Such technologies may be able to rescue protective immune responses to antigens that have previously failed by conventional bolus immunization, if immunodominance of non-neutralizing epitopes was a factor in their failure.

## AUTHOR CONTRIBUTIONS

K.M.C. performed AIM assays, ELISAs and BCR expression assays and analyzed flow cytometry, ELISA and BCR sequence data. D.G.C., C.A.E., E.H.G., Y.C., and F.V. performed NHP experiments and Env-specific B cell stains. A.A.U., V.K., and B.M. performed and analyzed BCR sequence and lineage analyses. C. N. and M.B.M. performed ELISAs. M.P. and R.B. performed and analyzed neutralization assays. A.N.W. performed single cell RNA-seq. C.A.C., B.N., and A.B.W. performed and analyzed EM experiments. S.R. performed CD38-CD71 GC B validation experiments. J.T.M. performed histology for antigen tracking study. S.C. designed the genome sequencing study. W.G., O.L.R. and C.T.W. annotated the germline Ig loci. N.P. and S.E.B. provided tissue for genomic sequencing. T.T. and D.J.I. provided the ISCOMs-type adjuvant. S.M., D.W.K., W.R.S. provided immunogens and Env probes. K.M.C. and S.C. prepared the manuscript, with input from other authors. D.J.I. and S.C. conceived of the study. S.C. and G.S. supervised the study.

## ACKNOWLEDGEMENTS

We thank Chai Fungtammasan and Brett Hannigan of DNAnexus for assembly of the rhesus macaque genome and helpful discussions, and Sanjeev Gumber of Yerkes National Primate Research Center for assistance with tissue isolation for genomic sequencing. This work was funded by NIH NIAID grant R01 AI125068 (D.J.I. and S.C.), NIH NIAID 1UM1Al100663 to the Scripps CHAVI-ID (SC, GS, DJI, WS, ABW, DRB), National Primate Research Center funding (P51 RR000165/OD011132 to the Yerkes National Primate Research Center) and NIH NIAID UM1AI124436 to the Emory Consortium for Innovative AIDS Research. C.A.C. is supported by NIH F31 Ruth L. Kirschstein Predoctoral Award Al131873 and by the Achievement Rewards for College Scientists Foundation.

## STAR Methods

### CONTACT FOR REAGENT AND RESOURCE SHARING

Further information and requests for resources and reagents should be directed to and will be fulfilled by the Lead Contact, Shane Crotty.

### EXPERIMENTAL MODEL AND SUBJECT DETAILS

#### Rhesus Macaques

Outbred Indian RMs (*Macaca mulatta*) were sourced and housed at the Yerkes National Primate Research Center and maintained in accordance with NIH guidelines. This study was approved by the Emory University Institutional Animal Care and Use Committee (IACUC). When osmotic pumps were implanted, animals were kept in single, protected contact housing. At all other times, animals were kept in paired housing. Animals were treated with anesthesia and analgesics for procedures as per veterinarian recommendations and IACUC approved protocols. In all studies, animals were grouped to divide age, weight and gender as evenly as possible.

Osmotic pump study: Animals were between 2.5 – 3 years of age at time of 1^st^ immunization. Bolus group 2: 2 males (M), 1 female (F); 2w pump group: 3M, 1 F; 4w pump group: 3M, 1 F.

Dose escalation study: Animals were between 3 – 6.5 years of age at time of 1^st^ immunization. Bolus group 1: 3M, 3F; escalating dose group: 2M, 4F.

Antigen tracking study: animals were between 3 – 6 years of age at time of immunization. Bolus group: 2M, 1F; pump group: 3M; escalating dose group: 3M.

### METHOD DETAILS

#### Immunizations

Osmotic pump study: Animals were immunized at 2 time points: week 0 and week 8. All immunizations were administered subcutaneously (SubQ) divided between the left and right mid-thighs. Bolus animals were given two SubQ injections of 50μg of Olio6-CD4ko + 187.5 units (U) of saponin adjuvant in PBS, for a total of 100μg Olio6-CD4ko trimer protein + 375U of saponin adjuvant. At week 0, osmotic pumps (Alzet, models −2002 and −2004) were loaded with 50μg Olio6-CD4ko + 187.5U saponin adjuvant, for a total of 100μg Olio6-CD4ko trimer + 375U of saponin adjuvant. Pumps were implanted SubQ in the same location as bolus immunizations. At week 8, osmotic pump animals were immunized with osmotic pumps loaded each with 25μg Olio6-CD4ko + 93.75U saponin adjuvant. At the end of the osmotic pump delivery, a SubQ bolus immunization of 25μg Olio5-CD4ko + 93.75U was given in each leg, totaling 50μg Olio6-CD4ko + 187.5U saponin adjuvant at weeks 12 and 14 for 2 week and 4 week osmotic pump groups, respectively.

Dose escalation study: Animals were immunized at 3 time points: weeks 0, 10, and 24. All immunizations were administered SubQ in the left and right mid-thighs. Bolus animals were given two injections of 50μg of Olio6 + 187.5U of saponin adjuvant in PBS, for total of 100μg immunogen and 375U saponin adjuvant at weeks 0 and 8. At week 24, two injections of 150μg of Olio6 + 187.5U saponin adjuvant were administered for a total of 300μg Olio6 + 375U saponin adjuvant. For each immunization, escalating dose animals were given seven injections of Olio6 and saponin adjuvant in each thigh over 12 days (on days 0, 2, 4, 6, 8, 10, 12 for each immunization). The total doses of Olio6 at each injection during the first two immunizations were: 0.2, 0.43, 1.16, 3.15, 8.56, 23.3, 63.2μg (the doses per immunization site were 0.1, 0.215, 0.58, 1.575, 4.28, 11.65, 31.6μg). The total doses of Olio6 at each injection during the third immunization were: 0.6, 1.29, 3.48, 9.45, 25.68, 69.9, 189.6μg (the doses per immunization site were 0.3, 0.645, 1.74, 4.725, 12.84, 34.95, 94.8μg). The total doses of saponin adjuvant at each injection during all immunizations were: 0.75, 1.61, 4.35, 11.81, 32.1, 87.38, 237.0U (the doses per immunization site were 0.375, 0.805, 2.175, 5.905, 16.05, 43.69, 118.5U).

Antigen tracking study: Animals were immunized at week 0 with a total dose of 100ug untagged MD39 conjugated to Alexa Fluor 647. All immunizations were administered SubQ in the left and right mid-thighs. Bolus animals were given 2 injections of 50μg MD39 + 187.5U of saponin adjuvant in PBS. Osmotic pumps (Alzet, models 2002) were loaded with 50μg MD39 + 187.5U saponin adjuvant. Escalating dose animals were given a series of 7 injections over 12 days (on days 0, 2, 4, 6, 8, 10, 12 for each immunization). The total dose of MD39 at each injection were: 0.2, 0.43, 1.16, 3.15, 8.56, 23.3, 63.2μg (the doses per immunization site were 0.1, 0.215, 0.58, 1.575, 4.28, 11.65, 31.6μg). The total doses of saponin adjuvant at each injection were: 0.75, 1.61, 4.35, 11.81, 32.1, 87.38, 237.0U (the doses per immunization site were 0.375, 0.805, 2.175, 5.905, 16.05, 43.69, 118.5U). Animals were sacrificed at 2 days after immunization (bolus, d2; pumps, d16; escalating dose, d14). All inguinal LNs were harvested and fixed in PLP buffer (pH7.4 50mM PBS + 100mM lysine, 1% paraformaldehyde, 2mg/mL sodium periodate) for 1 week at 4°C and then washed and stored in PBS with 0.05% sodium azide at 4°C until used for imaging.

#### Lymph node fine needle aspirates, whole LN biopsy tissue, blood collection and processing

LN FNAs were used to sample at both right and left inguinal LNs. FNAs were performed by a veterinarian. Draining lymph nodes were identified by palpitation. Cells were collected by passing a 22-gauge needle attached to a 3mL syringe into the lymph node 4 times. Samples were expelled into RPMI containing 10% fetal bovine serum, 1X penicillin/streptomycin. Samples were centrifuged and Ammonium-Chloride-Potassium (ACK) lysing buffer was used if sample was contaminated with red blood cells. Excisional LNs were conducted at weeks 12 (bolus) or 14 (osmotic pump groups). LNs were dissociated through 70μM strainers and washed with PBS. Blood was collected at various time points into CPT tubes for PBMC and plasma isolation. Serum was isolated using serum collection tubes and frozen.

#### ISCOMs-class saponin adjuvant

The adjuvant used for all the described studies was a ISCOM-like saponin nanoparticle comprised of self-assembled cholesterol phospholipid, and Quillaja saponin prepared as previously described (Lövgren-Bengtsson and Morein, 2000). Briefly, 10 mg each of cholesterol (Avanti Polar Lipids) and DPPC (Avanti Polar Lipids) were dissolved separately in 20% MEGA-10 (Sigma-Aldrich) detergent at a final concentration of 20 mg/mL and 50 mg Quil-A saponin (InvivoGen) was dissolved in MilliQ H_2_O at a final concentration of 100 mg/mL. Next, DPPC solution was added to cholesterol followed by addition of Quil-A saponin in rapid succession and the volume was brought up with PBS for a final concentration of 1 mg/mL cholesterol and 2% MEGA-10. The solution was allowed to equilibrate at 25°C overnight, followed by 5 days of dialysis against PBS using a 10k MWCO membrane. The adjuvant solution was filter sterilized using a 0.2 μm Supor syringe filter, concentrated using 50k MWCO centricon filters, and further purified by FPLC using a Sephacryl S-500 HR size exclusion column. Each adjuvant batch was finally characterized by negative stain transmission electron microscopy (TEM) and dynamic light scattering (DLS) to confirm uniform morphology and size and validated for low endotoxin content by Limulus Amebocyte Lysate assay (Lonza). Final adjuvant concentration was determined by cholesterol quantification (Sigma-Aldrich).

#### Immunogen and probe generation

Olio6, Olio6-CD4ko, and MD39 were generated as previously described(Kulp et al., 2017). Avi-tagged Olio6, Olio6-CD4ko, and MD39 DNA constructs were synthesized, protein was produced and purified, and the proteins were then biotinylated using BirA-500 (Avidity) and assessed for biotin conjugation efficiency using SDS-PAGE. All Env immunogens and probes contained a six histidine tag (His tag) for purification. Immunogens were tested for endotoxin contamination with Endosafe PTS (Charles River). Proteins with an endotoxin level <10 EU/mg were used in immunizations. Immunogens and probes were aliquoted and kept frozen at −80°C until immediately before use.

#### Flow cytometry and cellular analyses

Biotinylated protein were individually premixed with fluorochrome-conjugated streptavidin (SA-Alexa Fluor 647 or SA-Brilliant Violet 421) at RT for 20 minutes. Olio6-CD4ko probes were used in figures 1, 3, 6, and S1 (osmotic pump study) from weeks −1 to 8. Olio6 probes were used from weeks 9 to 14. Olio6 and Olio6-CD4ko differ by a single amino acid (Kulp et al., 2017). MD39 probes were used in figures 8 and S8 (dose escalation study). MD39 is closely related to Olio6.

For the full LN GC panel, cells were incubated with probes for 30 minutes at 4°C, washed twice and then incubated with surface antibodies for 30 minutes at 4°C. Cells were fixed and permeabilized for 30 minutes using FoxP3/Transcription Factor Staining Buffer Set (Thermo Scientific) according to manufacturer’s protocols. Cells were stained with intranuclear antibodies in 1X permeabilization buffer for 30 minutes, 4°C. Cells were washed twice with 1x permeabilization buffer and acquired on an LSR I (BD Biosciences). For Ag-specific B cell sort panels, cells were incubated with probes for 30 minutes at 4°C, washed twice and then incubated with surface antibodies for 30 minutes at 4°C. Cells were sorted on a FACSAria II.

For the osmotic pump study, full LN GC panel was used on fresh cells at weeks −2, 1–7, 9–12, 14. At weeks 7,12, and 14, cells were sorted using the Ag-specific B cell sort panel. Cells were stained fresh at week 7 and single cell sorted. At weeks 12 (bolus) and week 14 (osmotic pump animals), biopsied LNs were thawed, stained and bulk sorted for BR sequencing. Sorted cells were defined as Viability dye^−^ CD4^−^ CD8a^−^ CD16^−^ CD20^+^ (IgM^+^ IgG^+^)^−^ Olio6-Alexa647^+^ Olio6-BV421^+^. For the dose escalation study, the full LN GC panel was used at every time point. Data reported are raw flow cytometry values at each time point.

Validation of CD38 and CD71 as surface markers of GC B cells: frozen, biopsied mesenteric LNs were used. Cells were stained as described above.

B cell analysis: LN FNA samples 3% of the LN on average. Because of the nature of the technique, some samples do not have enough cells to be included in the analyses. Generally, for GC and Env-specific B cell gating, a threshold of 1,000 and 10,000 B cells, respectively, is used. For Env-specific GC B cell gating, a threshold of 1,000 GC B cells is used.

Inferred memory B cells: Memory B cells (% Env^+^ or Env^hi^) were calculated as the percentage of Env-specific or high-affinity Env-specific B cells that were not Bcl6^+^ Ki67^+^ or CD38^−^ CD71^+^. Memory Env^+^ and Env^hi^ (% B) cells were calculated as % Env^+^ (% B cells) - % Env^+^ GC B (% B) and % Env^hi^ (% B cells) - % Env^hi^ GC B (% B), respectively.

Area under the curve [AUC]: AUC was calculated for individual LNs. For figures 1 and S1, AUC was calculated from weeks 1, 3 to 7. Bolus gr1 did not have FNA data at week 1. For these samples, the median of the week 1 values from bolus gr2 was used. Raw values were used at other time points. For figure 2, AUC was calculated from weeks 1, 3 to 6. GC Tfh frequencies were not collected for bolus grp2, 2w pumps or 4w pump animals at week 7. For figure 3, AUC was calculated between weeks 9 and 12 because of poor cell recovery at weeks 8 and 14. For figures 8 and S8, AUC was calculated between weeks 3–7 (1^st^ immunization) and between weeks 11–15 (2^nd^ immunization) using raw values. Parameters used: baseline = 0; peaks less than 10% of distance from minimum to maximum y were ignored.

#### Antigen-specific CD4^+^ T cell assay

AIM assays were conducted as previously described (Dan et al., 2016; Havenar-Daughton et al., 2016b).

Osmotic pump study: Frozen macaque lymph nodes from week 12 (bolus animals) or week 14 (osmotic pump animals) were thawed. Cells were treated with DNAse (Stemcell Technologies) for 15 minutes, 37°C washed and then rested for 3 hours. Cells were cultured under the following conditions: media only (RPMI containing 10% fetal bovine serum, 1X penicillin/streptomycin, 2mM L-glutamine), 5ug/mL Olio6-CD4ko peptide megapool, or 1ng/mL SEB (positive control, Toxin Technology, Inc.). After 18 hours, cells were stained and acquired on FACSCelesta (BD Biosciences).

Dose escalation study: About 50% of lymphocytes are lost during the freeze-thaw process. To maximize the number of viable cells to identify Env-specific CD4^+^ cells, cells were shipped overnight at 4°C to LJI. Cells were centrifuged and treated with DNAse for 15 minutes, 37°C. Cells were washed, cultured for 18 hours under the conditions described above. All values reported are background subtracted ((% OX40^+^ 4-1BB^+^ CD4^+^ (Env-stimulated condition) – % OX40^+^ 4-1BB^+^ CD4^+^ (unstimulated condition)).

#### Whole genome sequencing and genome assembly

High molecular weight (>50kb) genomic DNA was isolated from the kidney of a perfused, female rhesus macaque. A full genome 30kb library was prepared according to manufacturer’s protocols. Sequencing was performed on a PacBio RS II (Pacific Biosciences). Genome assembly was performed using FALCON and FALCON-Unzip (Pacific Biosciences) (Chin et al., 2016). The final assembly contained 1633 contigs made up of 2.83 Gbp. The N50 contig length is 8.4Mbp, with a maximum contig length of 28.8Mbp.

#### Immunoglobulin loci annotation

Primary contigs from FALCON/FALCON-Unzip assemblies containing IG sequences were identified by aligning V, D, and J sequences from multiple sources, including sequences for RM and the crab-eating macaque (*Macaca fascicularis*) from the IMGT reference directory (http://www.imgt.org/vquest/refseqh.html) and (Corcoran et al., 2016) using BLAT (Kent, 2002). Gene annotation of primary contigs was carried out in two stages: (1) rough coordinates in each contig harboring putative V, D, and J segments were identified by mapping existing sequences (i.e., those noted above for contig identification, as well human IG D and J gene sequences from IMGT); followed by (2) manual curation, during which precise 5’ and 3’ gene segment boundaries were determined for each annotation, based on alignments to previously reported sequences, as well as the identification of flanking recombination signal sequence (RSS) heptamers within the contig assembly. Each gene annotation was assigned to a given subfamily based on the closest matching published sequence. Only ORF annotations lacking premature stop codons and/or insertion-deletions resulting in drastic frameshifts were considered.

Additional V gene allelic variants in the IGH, IGK, and IGL loci were identified by mapping PacBio raw reads back to IG-associated primary and alternate contigs from the FALCON/FALCON-Unzip assemblies using BLASR (Chaisson and Tesler, 2012). Putative heterozygous ORF genes were identified based on variants present in PacBio reads mapping to a given ORF locus (**Fig 6C**). To characterize putative alternate alleles, raw reads were partitioned and assembled locally at heterozygous ORFs using MsPAC (Rodriguez et al., *in prep*; https://bitbucket.org/oscarlr/mspac). Raw reads and assembled allelic variants were visually inspected in the context of primary and alternate FALCON/FALCON-Unzip contigs and confirmed using the Integrated Genomics Viewer (Robinson et al., 2011; Thorvaldsdóttir et al., 2013). To classify genes/alleles annotated from PacBio assembly data as “known” or “novel”, sequences were cross-referenced with the RM IMGT reference database and publicly available sequences in the NCBI nucleotide collection using BLAT and BLAST (https://blast.ncbi.nlm.nih.gov/Blast.cgi). respectively.

#### Bulk BCR sequencing

The protocol for rhesus macaque repertoire sequencing was obtained by courtesy of Dr. Daniel Douek, NIAID/VRC (Huang et al., 2016). Bulk Env-specific B cells were sorted into 350uL Qiagen RLT buffer. RNA was extracted using the RNeasy Micro-DNase Digest protocol (QIAGEN) on QIAcube automation platforms (Valencia, CA). Reverse transcription (RT) was performed using Clontech SMARTer cDNA template switching: 5’ CDS oligo(dT) (12 μM) was added to RNA and incubated at 72°C for 3 minutes and 4°C for at least 1 minute. The RT mastermix (5x RT Buffer (250 mM Tris-HCl (pH 8.3), 375 mM KCl, 30 mM MgCl_2_), Dithiothreitol, DTT(20 mM), dNTP Mix (10 mM). RNAse Out (40U/μL), SMARTer II A Oligo (12 μM), Superscript II RT (200U/μL)) was added to the reaction and incubated at 42°C for 90 minutes and 70°C for 10 minutes. First-strand cDNA was purified using AMPure XP beads (Beckman Coulter). Following RT, two PCR rounds were carried out to generate immunoglobulin amplicon libraries compatible with Illumina sequencing. All oligos were ordered from Integrated DNA Technologies. The first PCR amplification was carried out using KAPA Real-Time Library Amplification Kit (Kapa Biosciences). cDNA was combined with master mix (2X KAPA PCR Master Mix, 12 μM μL 5PIIA and 5 μL IgG/IgK/IgL Constant Primer (2 μM) (Francica et al., 2015)). The amplification was monitored using realtime PCR and was stopped during the exponential phase. The amplified products were again purified using AMPure XP beads. A second round of PCR amplification was carried out for addition of barcodes and Illumina adapter sequences: master mix (2X KAPA PCR Master Mix 2x, SYBR Green 1:10K, Nuclease-free water), 10 μM of P5_Seq BC_XX 5PIIA, 10 μM of P7_i7_XX IgG/IgK/IgL and were combined with amplified Immunoglobulin from the first round PCR and amplified using real-time PCR monitoring. The P5_Seq BC_XX 5PIIA primers contain a randomized stretch of four to eight random nucleotides. This was followed by purification with AMPure XP beads. A final PCR step was performed for addition of remaining Illumina adaptors by mixing master mix (2X KAPA PCR Master Mix, 10 μM P5_Graft P5_seq, Nuclease-free water), 10 μM of P7_ i7_XX IgG/IgK/IgL oligo and amplified products from the previous PCR step followed by purification with AMPure XP beads. The quality of library was assessed using Agilent Bioanalyzer. The amplicon libraries were pooled and sequenced on an Illumina MiSeq as a 309 paired-end run.

#### Single cell RNA-seq

Single cells were sorted by flow cytometry into 10 uL of QIAGEN RLT buffer. RNA was purified using RNACleanXP Solid Phase Reversible Immobilization (SPRI) beads (Beckman Coulter). Full-length cDNA amplification of single-cells was performed using a modified version of the SMART-Seq II protocol {Picelli 2014}, as described previously{Upadhyay 2018}. Amplified cDNA was fragmented using Illumina Nextera XT DNA Library Preparation kits and dual-indexed barcodes were added to each sample. Libraries were validated using an Agilent 4200 Tapestation, pooled, and sequenced at 101 SR on an Illumina HiSeq 3000 to an average depth of 1 M reads in the Yerkes NHP Genomics Core (http://www.yerkes.emory.edu/nhp_genomics_core/).

#### V gene and somatic hypermutation analyses

Illumina bcl files from IgG, IgK and IgL amplicons were converted to fastq files usng the bcl2fastq tool. FastQC v0.11.5 (Andrew, 2010) was used to check the quality of fastq files. The repertoire sequence analysis was carried out using the pRESTO 0.5.6, Change-O 0.3.12, Alakazam 0.2.10.999 and SHazaM 0.1.9 packages from the Immcantation pipeline (Gupta et al., 2015; Vander Heiden et al., 2014). Pre-processing was performed using tools in the pRESTO package. Paired-end reads were first assembled with AssemblePairs tool. Reads with a mean quality score of less than 20 were filtered out using FilterSeq. The MaskPrimers tool was used to remove the forward primers and the random nucleotides from the assembled sequences. Data from each of two technical replicates were combined. Duplicates were removed and the duplicate counts were obtained for each unique sequence using CollapseSeq. SplitSeq was used to select sequences that had duplicate counts of at least two to eliminate singletons that may arise due to sequencing errors. The pre-processed sequences were then annotated using IgBLAST v1.6.1(Ye et al., 2013).

Since the IMGT database (Lefranc and Lefranc, 2001) is lacking several V genes, a custom IgBLAST database was created for V genes using sequences from the genomic assembly in this study or by combining sequences from previously published studies (Corcoran et al., 2016; Lefranc and Lefranc, 2001; Ramesh et al., 2017; Sundling et al., 2012) and the sequences from the assembly in this study. The protein sequences for V genes from all these datasets were combined. The sequences were aligned using MUSCLE v3.8.1551 (Edgar, 2004) and only the V genes with complete sequence and no unknown amino acid (X) were selected. The corresponding nucleotide sequences of these V genes were clustered using CD-HIT v4.7 (Fu et al., 2012) to remove 100% redundant sequences. The protein sequences for this non-redundant set were submitted to the IMGT DomainGapAlign tool (Ehrenmann and Lefranc, 2011; Ehrenmann et al., 2010) to obtain gapped V sequences. Corresponding gaps were introduced in the nucleotide sequences and the positions for framework (FR) and complementarity-determining regions (CDR) regions determined using custom scripts. These sequences were used to create the IgBLAST database for V genes. The databases for J and D genes was obtained from the IgBLAST ftp site (ftp://ftp.ncbi.nih.gov/blast/executables/igblast/release/internal_data/rhesus_monkey/). The annotations from IgBLAST were saved into a Change-O database and functional sequences were selected using Change-O. The gene usage and clonal frequencies were obtained from the Alakazam package and SHM estimations were obtained from the SHazaM package.

To obtain paired heavy and light chain sequences from single cell RNA-Seq data, we used the BALDR pipeline, as previously described (Upadhyay et al., 2018), with the Unfiltered method for rhesus macaques.

#### Lineage analysis

For the quantification of B cell lineages, two independent analyses were performed with largely equivalent results.

Lineage analyses in figures 5 and S4 utilized only the sequences from the genomic assembly generated in this study. The annotations from IgBLAST were saved into a Change-O database and functional sequences were selected using Change-O. The functional sequences were assigned to a clone using a custom script based on the following criteria: (i) same V gene, (ii) same J gene, (iii) same CDR3 length and (iv) percentage identity of CDR3 nucleotide sequence > 85%. The analysis was also performed with the larger IgBLAST database with comparable results.

Phylogenetic trees were generated using the larger IgBLAST database described above. Lineage assignment was performed using a clustering procedure that exploited both germline inference and sequence similarity. Two sequences were deemed to potentially belong to the same lineage when: (i) their inferred UCA sequences (ignoring the junction and D region) are within 1% of each other (using a kmer-based distance approximation from (Kumar et al., 2018) for computational efficiency), tolerating calls to closely related V and J genes; and (ii) when the length-normalized Levenshtein distance between their junction+D sequences is within 10%. The clustering algorithm itself maintains a set of candidate lineages, storing all sequences for each lineage, and each new sequence in turn is added to the lineage where the largest proportion of sequences match the above two criteria. If no existing candidate cluster has >50% of its reads match the new sequence, then that sequence is used to seed a new candidate cluster containing this sequence as its sole member. Where members of a lineage had different inferred UCA sequences, the modal UCA was chosen as the UCA for the entire lineage. This lineage clustering algorithm was implemented in the Julia language for scientific computing (v0.6.2). Each lineage was aligned with MAFFT (Katoh and Standley, 2013), and phylogenetic trees were inferred using FastTree2 (Price et al., 2010). Phylogenies were visualized using FigTree (http://tree.bio.ed.ac.uk/software/figtree/). using automated coloring and annotation scripts implemented in Julia.

#### ELISAs

BG505 SOSIP, BG505 gp120 and His ELISAs: 96-well Maxisorp plates (Thermo Fisher Scientific) plates were coated with streptavidin at 2.5μg/mL (Thermo Fisher Scientific) overnight at 4C. Plates were washed with PBS + 0.05% Tween (PBS-T) three times. Biotinylated BG505, biotinylated His peptide conjugated to mouse CD1d or biotinylated gp120 was diluted to 1.0μg/mL in PBS + 1% BSA were captured for 2 hours, 37°C. Plates were washed three times and then blocked with PBS^+^ 3% BSA for 1 hour, RT. Plasma samples or monoclonal antibodies were serially diluted in PBS + 1% BSA and incubated for 1 hour, RT. Plates were washed three times and horseradish peroxidase goat anti-rhesus IgG (H + L) secondary (Southern Biotech) was added at 1:3000 dilution in PBS + 1% PBS for 1 hour, RT. Plates were washed three times with PBS-T and absorption was measured at 450nm following addition of TMB substrate (Thermo Scientific). We calculated endpoint titers for BG505 SOSIP and His peptide ELISAs using GraphPad Prism v7.0. Antibody data panels show geometric mean titers with geometric SD.

Lectin-capture BG505 trimer ELISA: To maximize access to the base of the trimer, we utilized a lectin-capture assay. Env trimer is heavily glycosylated, except at the base. Capture with a lectin, which binds glycans, increases the likelihood that the base will be exposed more than in a streptavidin-capture ELISA. Half-area 96- well high binding plates (Corning) were coated with 5μg/mL lectin from *Galanthus nivalis* (snowdrop) (Sigma) in PBS overnight at 4°C. Plates were washed with 0.05% PBS-Tween (PBS-T) three times. 1 ug/ml BG505 trimer in PBS + 1% BSA was bound to plates for 2 hours at 37°C and then washed three times. Plates were blocked with PBS + 3% BSA for 1 hour, RT. Monoclonal antibodies were serially diluted in PBS + 1% BSA and incubated for 1.5 hours at RT. Plates were washed three times and incubated with horseradish peroxidase goat anti-rhesus IgG (H+L) secondary antibody (Southern Biotech) at 1:3000 in PBS + 1% BSA for 1 hr, RT. Plates were washed five times with PBS-T and absorption was measured at 450nm following addition of TMB substrate (Thermo Fisher Scientific).

Cross-competition trimer ELISA: We used a modified lectin capture ELISA for this assay. Plates were coated with GNL and BG505 and blocked as previously described. Plates were incubated with 0 or 10μg/mL PGT121 or 19R (fab) in PBS + 1% BSA for 1.5 hours at RT. Plates were washed three times with PBS-T. 5μg/mL of BDA1 (whole antibody) was added for 1 hour at RT and then washed three times with PBS-T before incubation with horseradish peroxidase goat anti-human IgG, Fcg fragment specific (Jackson ImmunoResearch) at 1:5000 in PBS + 1% BSA for 1hr, RT. Plates were washed five times with PBS-T and absorption was measured at 450nm following addition of TMB substrate (Thermo Fisher Scientific).

#### 19R

The genes encoding the 19R rhesus macaque IgG1 heavy chain and kappa light chain were synthesized and separately cloned into the pcDNA3.4 plasmid by Thermo Fisher Scientific. The 19R IgG was expressed in Expi293 cells and purified using Protein A by Thermo Fisher Scientific. 19R Fab was generated by digesting 19R IgG using the Pierce Fab Preparation Kit (Thermo Fisher Scientific).

#### Pseudovirus neutralization assay

Neutralization assays were performed as previously described (Pauthner et al., 2017). Neutralization titers are reported as IC_50_ titers. All ELISA and neutralization Ab data panels show geometric mean titers with geometric SD.

#### Monoclonal EM analysis

The heavy and light chains of BDA1 (HC:QVQLQESGPGLVKPSETLSLTCAVSGASISIYWWGWIRQPPGKGLEWIGEIIGSSGSTNSNPSFKSRVTISKDASKNQFSLNLNSVTAADTAVYYCVRVGAAISLPFDYWGQGVLVTVSS, LC: SYELTQPPSVSVSPGQTARITCSGDALPKKYAYWFQQKPGQSPVLIIYEDNKRPSGIPERFSGSSSGTVATLTISGAQVEDEGDYYCYSRHSSGNHGLFGGGTRLTVL) were codon-optimized, synthesized and cloned into pFUSE2ss-CHIg-hG1 and pFUSE2ss-CLIg-hl2, respectively, by GenScript. Antibodies were expressed and purified by GenScript. Fab was generated using Pierce Fab preparation kit (Thermo Fisher Scientific). 15 μg of BG505 SOSIPv5.2 Env trimer (untagged) was complexed with 41μg BDa1 Fab at room temperature overnight in a total reaction volume of 50 μL. The complex was diluted 1:20 with TBS and 3 μL was applied to a glow-discharged, carbon-coated 400-mesh copper grid and blotted off after 15 seconds. 3 μL of 2% (w/v) uranyl formate stain was applied and immediately blotted off, followed by another application of 3 μL of stain for 45 seconds, blotted once more, and allowed to air-dry. Images were collected via Leginon (Potter et al., 1999) using an FEI Talos microscope (1.98 Å/pixel; 72,000× magnification; 25 e^−^/ Å^2^). Particles were picked from the raw images using DoG Picker (Voss et al., 2009). 2D classification, 3D sorting and refinement of the complex was conducted using RELION 3.0b0 (Nakane et al., 2018).

#### Polyclonal EM analysis

Plasma from week 10 (bolus), week 12 (2 week pumps) or week 14 (4 week pumps) was diluted 4X with PBS and incubated with protein A sepharose beads (GE Healthcare) overnight at 4C. Resin was washed 3X with PBS and eluted with 0.1M glycine pH2.5 and immediately neutralized with 1M Tris-HCL pH 8. Fabs were purified using Pierce Fab preparation kit (Thermo). Fab was generated using Pierce Fab Preparation Kit (Thermo Scientific). Reaction was incubated with protein A sepharose resin for 1 hour, RT. Fabs were buffer exchanged using Amicon ultra 0.5ml centrifugal filters (Millipore Sigma).

Upon buffer exchange into TBS, 0.5 to 0.8 mg of total Fab was incubated overnight with 10 μg BG505 trimers at RT in ~36 μL total volume. The formed complexes were then separated from unbound Fab via size exclusion chromatography (SEC) using Superose 6 Increase 10/300 column (GE Healthcare) equilibrated with TBS. The flow-through fractions containing the complexes were pooled and concentrated using 100 kDa cutoff centrifugal filters (EMD Millipore). The final trimer concentration was adjusted to approximately 0.04 mg/mL prior to application onto carbon-coated copper grids.

Complexes were applied to glow-discharged, carbon-coated 400-mesh copper grids, followed by applying 3 μL of 2% (w/v) uranyl formate stain that was immediately blotted off, and followed by application of another 3 μL of stain for 45–60 s, and blotted once more. Stained grids were allowed to air-dry and stored under ambient conditions until imaging. Images were collected via Leginon (Potter et al., 1999) using a Tecnai T12 electron microscopes operated at 120 kV; ×52,000 magnification; 2.05 Å/pixel. In all cases, the electron dose was 25 e^−^/ Å^2^. Particles were picked from the raw images using DoG Picker (Voss et al., 2009) and placed into stacks using Appion software (Lander et al., 2009). 2D reference-free alignment was performed using iterative MSA/MRA) (Sorzano et al., 2010). Finally, the particle stacks were then converted from IMAGIC to RELION-formatted MRC stacks and subjected to RELION 2.1 2D and 3D classification (Scheres, 2012).

#### Histology

Selected LNs were embedded in 3% low melting temperature agarose (Sigma-Aldrich), and then sliced into 350 μm-thick sections using a vibratome. The slices were blocked and permeabilized for 2 days in PBS with 10% goat serum and 0.2% Triton-X-100, followed by staining for 3 days with BV421-labeled mouse anti-human CD35 clone E11 (BD Biosciences) and Alexa Fluor 488-labeled mouse anti-Ki67 clone B56 (BD Biosciences) in the blocking buffer. Stained slices were then washed for 3 days with PBS containing 0.2% Tween-20, and then mounted onto glass slides with coverslips. Imaging was performed on either a Leica SP8 or an Olympus FV1200 laser scanning confocal microscope using 10x objectives. Images were analyzed using ImageJ.

### QUANTIFICATION AND STATISTICAL ANALYSIS

Graphpad Prism 7.0 was used for all statistical analyses. Significance of differences in neutralization, BG505 binding titers, cellular frequencies and geoMFI were calculated using unpaired, two-tailed Mann-Whitney U tests. Differences in mutation frequencies between groups were calculated using unpaired Student’s t tests. Significance of differences in V gene use between groups were calculated using multiple t tests, corrected for multiple comparisons with a false discovery rate (FDR) of 5% (Benjamini, Krieger, and Yekutieli). Differences in BCR expression of GC vs non-GC B cells were calculated using paired, Wilcoxon test. Correlations between neutralization and cell frequencies were calculated using log transformed Ab titer values in two-tailed Pearson correlation tests.

### DATA AND SOFTWARE AVAILABILITY

The rhesus macaque germline Ig V, D and J reference genes and Env-specific B cell BCR sequences used in this paper are available at NCBI Sequence Read Archive (https://www.ncbi.nlm.nih.gov/sra). 3D EM reconstructions have been deposited in the Electron Microscopy Databank (http://www.emdatabank.org/) under the accession numbers listed in the Key Resources Table.

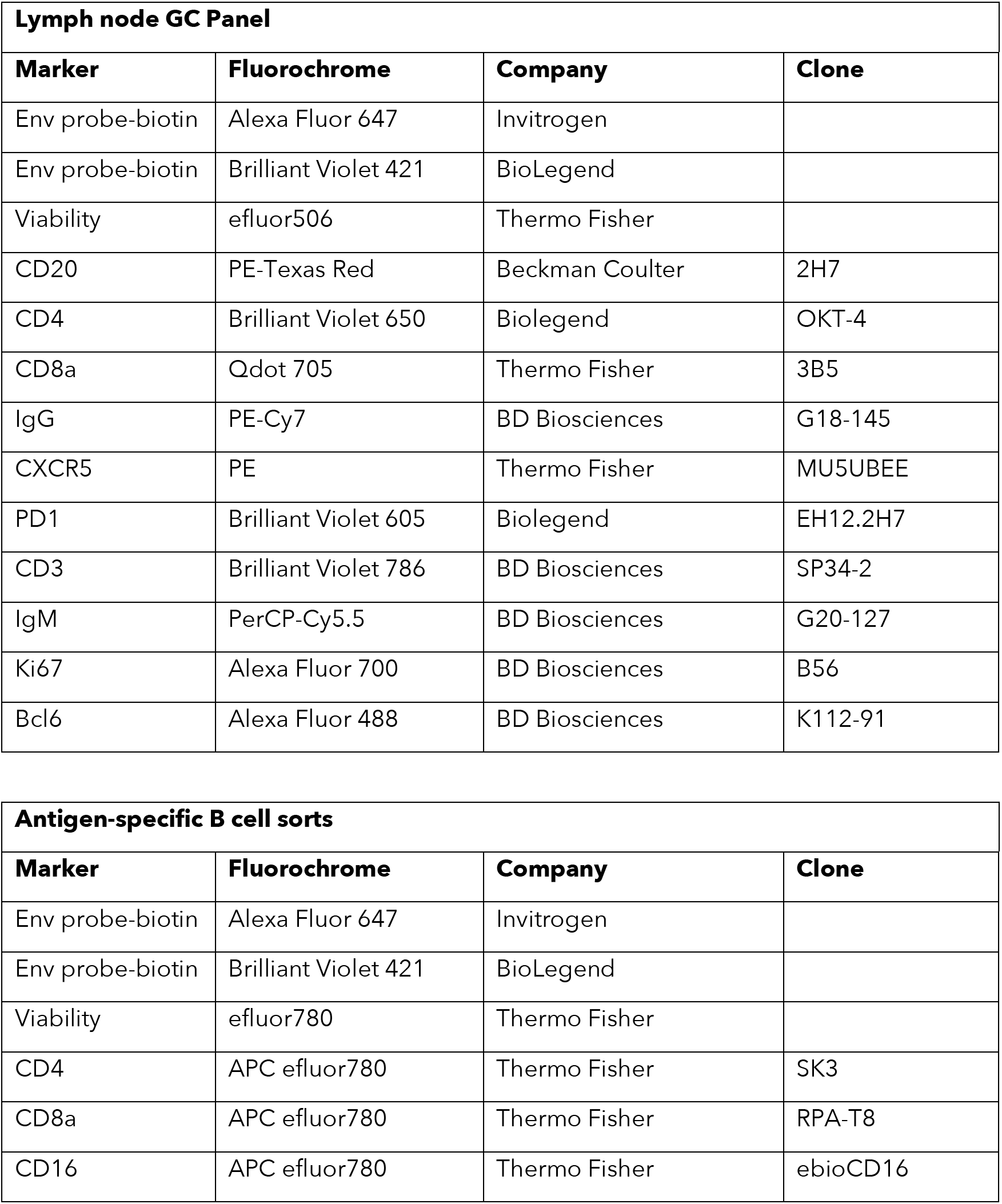

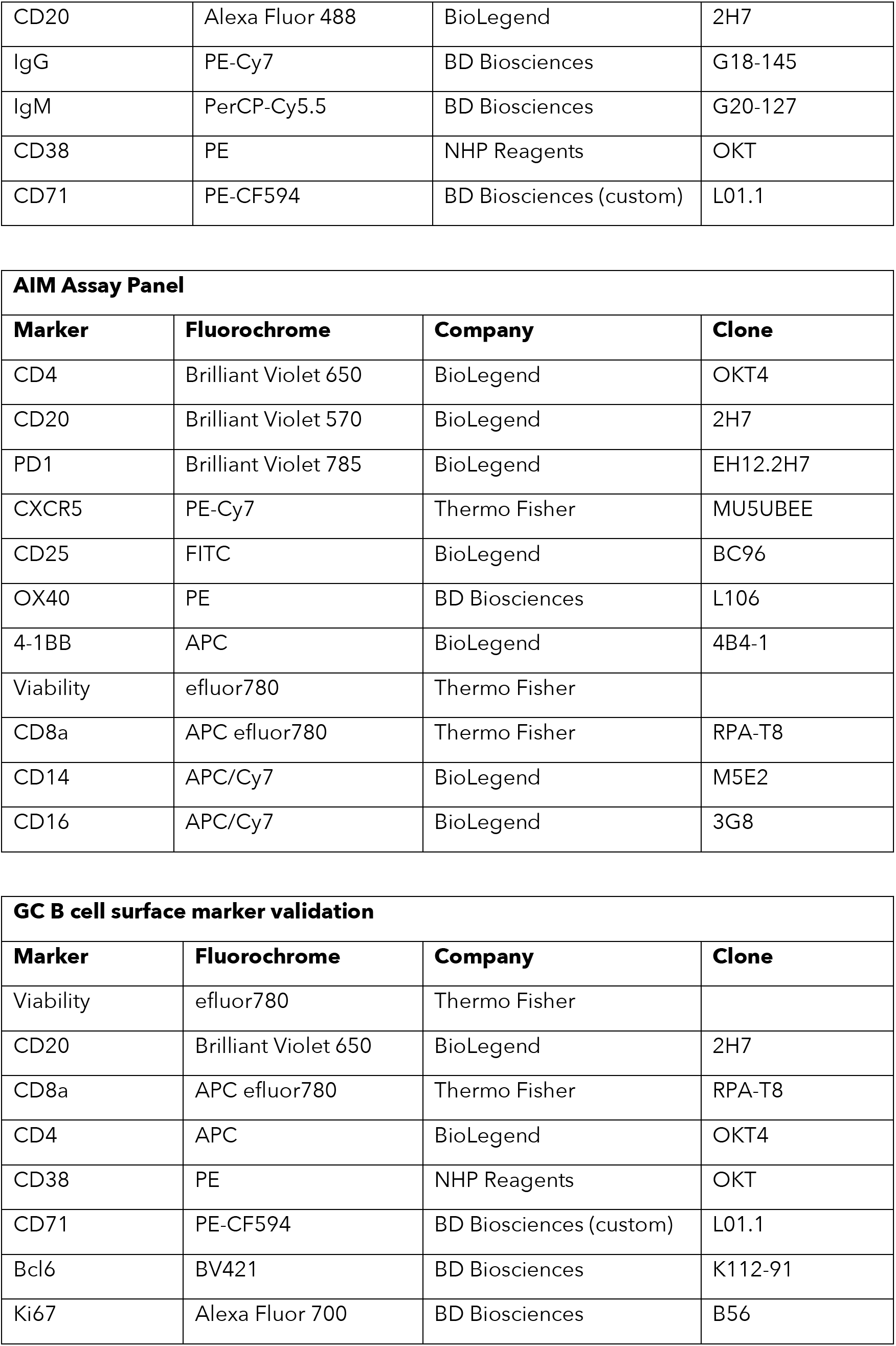

| Macaque BCR expression |  |  |  |
| --- | --- | --- | --- |
| Marker | Fluorochrome | Company | Clone |
| Viability | efluor780 | Thermo Fisher |  |
| CD4 | APC efluor780 | Thermo Fisher | SK3 |
| CD8a | APC efluor780 | Thermo Fisher | RPA-T8 |
| CD16 | APC/Cy7 | BioLegend | 3G8 |
| CD20 | Brilliant Violet 650 | BioLegend | 2H7 |
| Bcl6 | Alexa Fluor 647 | BD Biosciences | K112-91 |
| Ki67 | Alexa Fluor 700 | BD Biosciences | B56 |
| IgM | BV421 | BD Biosciences | G20-127 |
| IgG | PE | BD Biosciences | G18-145 |
| IgD | Alexa Fluor 488 | Southern Biotech |  |
| Lambda | Biotin | Miltenyi | IS7-24C7 |
| Streptavidin | Brilliant Violet 711 | BioLegend |  |

## REFERENCES

Abbott, R.K., Lee, J.H., Menis, S., Skog, P., Rossi, M., Ota, T., Kulp, D.W., Bhullar, D., Kalyuzhniy, O., Havenar-Daughton, C., et al. (2018). Precursor Frequency and Affinity Determine B Cell Competitive Fitness in Germinal Centers, Tested with Germline-Targeting HIV Vaccine Immunogens. Immunity 48, 133–146.e136.

Alkan, C., Sajjadian, S., and Eichler, E.E. (2011). Limitations of next-generation genome sequence assembly. Nat. Methods 8, 61–65.

Andrew, S. (2010). FastQC: A quality control tool for high throughput sequence data. 2010.

Andrews, S.F., Graham, B.S., Mascola, J.R., and McDermott, A.B. (2018). Is It Possible to Develop a “Universal” Influenza Virus Vaccine? Immunogenetic Considerations Underlying B-Cell Biology in the Development of a Pan-Subtype Influenza A Vaccine Targeting the Hemagglutinin Stem. Cold Spring Harb Perspect Biol 10, a029413.

Angeletti, D., and Yewdell, J.W. (2018). Understanding and Manipulating Viral Immunity: Antibody Immunodominance Enters Center Stage. Trends Immunol. 39, 549–561.

Angeletti, D., Gibbs, J.S., Angel, M., Kosik, I., Hickman, H.D., Frank, G.M., Das, S.R., Wheatley, A.K., Prabhakaran, M., Leggat, D.J., et al. (2017). Defining B cell immunodominance to viruses. Nat. Immunol. 18, 456–463.

Baiyegunhi, O., Ndlovu, B., Ogunshola, F., Ismail, N., Walker, B.D., Ndung’u, T., and Ndhlovu, Z.M. (2018). Frequencies of Circulating Th1-Biased T Follicular Helper Cells in Acute HIV-1 Infection Correlate with the Development of HIV-Specific Antibody Responses, and Lower Set Point Viral Load. J. Virol. 92, 2209.

Bianchi, M., Turner, H.L., Nogal, B., Cottrell, C.A., Oyen, D., Pauthner, M., Bastidas, R., Nedellec, R., McCoy, L.E., Wilson, I.A., et al. (2018). Electron-Microscopy-Based Epitope Mapping Defines Specificities of Polyclonal Antibodies Elicited during HIV-1 BG505 Envelope Trimer Immunization. Immunity 49, 288–300.e288.

Burton, D.R., and Hangartner, L. (2016). Broadly Neutralizing Antibodies to HIV and Their Role in Vaccine Design. Annu. Rev. Immunol. 34, 635–659.

Chaisson, M.J., and Tesler, G. (2012). Mapping single molecule sequencing reads using basic local alignment with successive refinement (BLASR): application and theory. BMC Bioinformatics 13, 238.

Chin, C.-S., Peluso, P., Sedlazeck, F.J., Nattestad, M., Concepcion, G.T., Clum, A., Dunn, C., O’Malley, R., Figueroa-Balderas, R., Morales-Cruz, A., et al. (2016). Phased diploid genome assembly with single-molecule real-time sequencing. Nat. Methods 13, 1050–1054.

Chowdhury, A., Del Rio, P.M.E., Tharp, G.K., Trible, R.P., Amara, R.R., Chahroudi, A., Reyes-Teran, G., Bosinger, S.E., and Silvestri, G. (2015). Decreased T Follicular Regulatory Cell/T Follicular Helper Cell (TFH) in Simian Immunodeficiency Virus-Infected Rhesus Macaques May Contribute to Accumulation of TFH in Chronic Infection. J. Immunol. 195, 3237–3247.

Cirelli, K.M., and Crotty, S. (2017). Germinal center enhancement by extended antigen availability. Curr. Opin. Immunol. 47, 64–69.

Corcoran, M.M., Phad, G.E., Vázquez Bernat, N., Stahl-Hennig, C., Sumida, N., Persson, M.A.A., Martin, M., and Karlsson Hedestam, G.B. (2016). Production of individualized V gene databases reveals high levels of immunoglobulin genetic diversity. Nat Commun 7, 13642.

Crotty, S. (2014). T follicular helper cell differentiation, function, and roles in disease. Immunity 41, 529–542.

Dan, J.M., Lindestam Arlehamn, C.S., Weiskopf, D., da Silva Antunes, R., Havenar-Daughton, C., Reiss, S.M., Brigger, M., Bothwell, M., Sette, A., and Crotty, S. (2016). A Cytokine-Independent Approach To Identify Antigen-Specific Human Germinal Center T Follicular Helper Cells and Rare Antigen-Specific CD4+ T Cells in Blood. J. Immunol. 197, 983–993.

DeMuth, P.C., Li, A.V., Abbink, P., Liu, J., Li, H., Stanley, K.A., Smith, K.M., Lavine, C.L., Seaman, M.S., Kramer, J.A., et al. (2013). Vaccine delivery with microneedle skin patches in nonhuman primates. Nat. Biotechnol. 31, 1082–1085.

DeMuth, P.C., Min, Y., Irvine, D.J., and Hammond, P.T. (2014). Implantable silk composite microneedles for programmable vaccine release kinetics and enhanced immunogenicity in transcutaneous immunization. Adv Healthc Mater 3, 47–58.

Duan, H., Chen, X., Boyington, J.C., Cheng, C., Zhang, Y., Jafari, A.J., Stephens, T., Tsybovsky, Y., Kalyuzhniy, O., Zhao, P., et al. (2018). Glycan Masking Focuses Immune Responses to the HIV-1 CD4-Binding Site and Enhances Elicitation of VRC01-Class Precursor Antibodies. Immunity 49, 301–311.e305.

Edgar, R.C. (2004). MUSCLE: multiple sequence alignment with high accuracy and high throughput. Nucleic Acids Res. 32, 1792–1797.

Ehrenmann, F., and Lefranc, M.-P. (2011). IMGT/DomainGapAlign: IMGT standardized analysis of amino acid sequences of variable, constant, and groove domains (IG, TR, MH, IgSF, MhSF). Cold Spring Harb Protoc 2011, 737–749.

Ehrenmann, F., Kaas, Q., and Lefranc, M.-P. (2010). IMGT/3Dstructure-DB and IMGT/DomainGapAlign: a database and a tool for immunoglobulins or antibodies, T cell receptors, MHC, IgSF and MhcSF. Nucleic Acids Res. 38, D301–D307.

Feng, Y., Tran, K., Bale, S., Kumar, S., Guenaga, J., Wilson, R., de Val, N., Arendt, H., DeStefano, J., Ward, A.B., et al. (2016). Thermostability of Well-Ordered HIV Spikes Correlates with the Elicitation of Autologous Tier 2 Neutralizing Antibodies. PLoS Pathog. 12, e1005767.

Francica, J.R., Sheng, Z., Zhang, Z., Nishimura, Y., Shingai, M., Ramesh, A., Keele, B.F., Schmidt, S.D., Flynn, B.J., Darko, S., et al. (2015). Analysis of immunoglobulin transcripts and hypermutation following SHIV(AD8) infection and protein-plus-adjuvant immunization. Nat Commun 6, 6565.

Fu, L., Niu, B., Zhu, Z., Wu, S., and Li, W. (2012). CD-HIT: accelerated for clustering the next-generation sequencing data. Bioinformatics 28, 3150–3152.

Gibbs, R.A., Rogers, J., Katze, M.G., Bumgarner, R., Weinstock, G.M., Mardis, E.R., Remington, K.A., Strausberg, R.L., Venter, J.C., Wilson, R.K., et al. (2007). Evolutionary and biomedical insights from the rhesus macaque genome. Science 316, 222–234.

Gitlin, A.D., Mayer, C.T., Oliveira, T.Y., Shulman, Z., Jones, M.J.K., Koren, A., and Nussenzweig, M.C. (2015). HUMORAL IMMUNITY. T cell help controls the speed of the cell cycle in germinal center B cells. Science 349, 643–646.

Gitlin, A.D., Shulman, Z., and Nussenzweig, M.C. (2014). Clonal selection in the germinal centre by regulated proliferation and hypermutation. Nature 509, 637–640.

Gupta, N.T., Vander Heiden, J.A., Uduman, M., Gadala-Maria, D., Yaari, G., and Kleinstein, S.H. (2015). Change-O: a toolkit for analyzing large-scale B cell immunoglobulin repertoire sequencing data. Bioinformatics 31, 3356–3358.

Havenar-Daughton, C., Abbott, R.K., Schief, W.R., and Crotty, S. (2018). When designing vaccines, consider the starting material: the human B cell repertoire. Curr. Opin. Immunol. 53, 209–216.

Havenar-Daughton, C., Carnathan, D.G., Torrents de la Peña, A., Pauthner, M., Briney, B., Reiss, S.M., Wood, J.S., Kaushik, K., van Gils, M.J., Rosales, S.L., et al. (2016a). Direct Probing of Germinal Center Responses Reveals Immunological Features and Bottlenecks for Neutralizing Antibody Responses to HIV Env Trimer. Cell Rep 17, 2195–2209.

Havenar-Daughton, C., Lee, J.H., and Crotty, S. (2017). Tfh cells and HIV bnAbs, an immunodominance model of the HIV neutralizing antibody generation problem. Immunological Reviews 275, 49–61.

Havenar-Daughton, C., Reiss, S.M., Carnathan, D.G., Wu, J.E., Kendric, K., Torrents de la Peña, A., Kasturi, S.P., Dan, J.M., Bothwell, M., Sanders, R.W., et al. (2016b). Cytokine-Independent Detection of Antigen-Specific Germinal Center T Follicular Helper Cells in Immunized Nonhuman Primates Using a Live Cell Activation-Induced Marker Technique. J. Immunol. 197, 994–1002.

Haynes, B.F., Gilbert, P.B., McElrath, M.J., Zolla-Pazner, S., Tomaras, G.D., Alam, S.M., Evans, D.T., Montefiori, D.C., Karnasuta, C., Sutthent, R., et al. (2012). Immune-correlates analysis of an HIV-1 vaccine efficacy trial. N. Engl. J. Med. 366, 1275–1286.

Hogenesch, H. (2002). Mechanisms of stimulation of the immune response by aluminum adjuvants. Vaccine 20, S34–S39.

Hogenesch, H. (2012). Mechanism of immunopotentiation and safety of aluminum adjuvants. Front Immunol 3, 406.

Hu, J.K., Crampton, J.C., Cupo, A., Ketas, T., van Gils, M.J., Sliepen, K., de Taeye, S.W., Sok, D., Ozorowski, G., Deresa, I., et al. (2015). Murine Antibody Responses to Cleaved Soluble HIV-1 Envelope Trimers Are Highly Restricted in Specificity. J. Virol. 89, 10383–10398.

Huang, J., Kang, B.H., Ishida, E., Zhou, T., Griesman, T., Sheng, Z., Wu, F., Doria-Rose, N.A., Zhang, B., McKee, K., et al. (2016). Identification of a CD4-Binding-Site Antibody to HIV that Evolved Near-Pan Neutralization Breadth. Immunity 45, 1108–1121.

Hutchison, S., Benson, R.A., Gibson, V.B., Pollock, A.H., Garside, P., and Brewer, J.M. (2012). Antigen depot is not required for alum adjuvanticity. Faseb J. 26, 1272–1279.

Jardine, J.G., Kulp, D.W., Havenar-Daughton, C., Sarkar, A., Briney, B., Sok, D., Sesterhenn, F., Ereño-Orbea, J., Kalyuzhniy, O., Deresa, I., et al. (2016). HIV-1 broadly neutralizing antibody precursor B cells revealed by germline-targeting immunogen. Science 351, 1458–1463.

Julien, J.-P., Cupo, A., Sok, D., Stanfield, R.L., Lyumkis, D., Deller, M.C., Klasse, P.J., Burton, D.R., Sanders, R.W., Moore, J.P., et al. (2013). Crystal structure of a soluble cleaved HIV-1 envelope trimer. Science 342, 1477–1483.

Katoh, K., and Standley, D.M. (2013). MAFFT Multiple Sequence Alignment Software Version 7: Improvements in Performance and Usability. Molecular Biology and Evolution 30, 772–780.

Kent, W.J. (2002). BLAT--the BLAST-like alignment tool. Genome Res. 12, 656–664.

Klasse, P.J., Ketas, T.J., Cottrell, C.A., Ozorowski, G., Debnath, G., Camara, D., Francomano, E., Pugach, P., Ringe, R.P., LaBranche, C.C., et al. (2018). Epitopes for neutralizing antibodies induced by HIV-1 envelope glycoprotein BG505 SOSIP trimers in rabbits and macaques. PLoS Pathog. 14, e1006913.

Klein, F., Mouquet, H., Dosenovic, P., Scheid, J.F., Scharf, L., and Nussenzweig, M.C. (2013). Antibodies in HIV-1 vaccine development and therapy. Science 341, 1199–1204.

Kong, R., Xu, K., Zhou, T., Acharya, P., Lemmin, T., Liu, K., Ozorowski, G., Soto, C., Taft, J.D., Bailer, R.T., et al. (2016). Fusion peptide of HIV-1 as a site of vulnerability to neutralizing antibody. Science 352, 828–833.

Kulp, D.W., Steichen, J.M., Pauthner, M., Hu, X., Schiffner, T., Liguori, A., Cottrell, C.A., Havenar-Daughton, C., Ozorowski, G., Georgeson, E., et al. (2017). Structure-based design of native-like HIV-1 envelope trimers to silence non-neutralizing epitopes and eliminate CD4 binding. Nat Commun 8, 1655.

Kumar, V., Vollbrecht, T., Chernyshev, M., Mohan, S., Hanst, B., Bavafa, N., Lorenzo, A., Ketteringham, R., Eren, K., Golden, M., et al. (2018). Long-read amplicon denoising. 1–10.

Kuraoka, M., Schmidt, A.G., Nojima, T., Feng, F., Watanabe, A., Kitamura, D., Harrison, S.C., Kepler, T.B., and Kelsoe, G. (2016). Complex Antigens Drive Permissive Clonal Selection in Germinal Centers. Immunity 44, 542–552.

Lander, G.C., Stagg, S.M., Voss, N.R., Cheng, A., Fellmann, D., Pulokas, J., Yoshioka, C., Irving, C., Mulder, A., Lau, P.-W., et al. (2009). Appion: an integrated, database-driven pipeline to facilitate EM image processing. J. Struct. Biol. 166, 95–102.

Lefranc, M.P., and Lefranc, G. (2001). The immunoglobulin factsbook.

Locci, M., Havenar-Daughton, C., Landais, E., Wu, J., Kroenke, M.A., Arlehamn, C.L., Su, L.F., Cubas, R., Davis, M.M., Sette, A., et al. (2013). Human circulating PD-1+CXCR3-CXCR5+ memory Tfh cells are highly functional and correlate with broadly neutralizing HIV antibody responses. Immunity 39, 758–769.

Lövgren-Bengtsson, K., and Morein, B. (2000). The ISCOM™ Technology. In Vaccine Adjuvants, (New Jersey: Humana Press), pp. 239–258.

Lyumkis, D., Julien, J.-P., de Val, N., Cupo, A., Potter, C.S., Klasse, P.J., Burton, D.R., Sanders, R.W., Moore, J.P., Carragher, B., et al. (2013). Cryo-EM structure of a fully glycosylated soluble cleaved HIV-1 envelope trimer. Science 342, 1484–1490.

Mascola, J.R., Snyder, S.W., Weislow, O.S., Belay, S.M., Belshe, R.B., Schwartz, D.H., Clements, M.L., Dolin, R., Graham, B.S., Gorse, G.J., et al. (1996). Immunization with envelope subunit vaccine products elicits neutralizing antibodies against laboratory-adapted but not primary isolates of human immunodeficiency virus type 1. The National Institute of Allergy and Infectious Diseases AIDS Vaccine Evaluation Group. J. Infect. Dis. 173, 340–348.

Mesin, L., Ersching, J., and Victora, G.D. (2016). Germinal Center B Cell Dynamics. Immunity 45, 471–482.

Montefiori, D.C., Roederer, M., Morris, L., and Seaman, M.S. (2018). Neutralization tiers of HIV-1. Curr Opin HIV AIDS 13, 128–136.

Moody, M.A., Pedroza-Pacheco, I., Vandergrift, N.A., Chui, C., Lloyd, K.E., Parks, R., Soderberg, K.A., Ogbe, A.T., Cohen, M.S., Liao, H.-X., et al. (2016). Immune perturbations in HIV-1-infected individuals who make broadly neutralizing antibodies. Science Immunology 1, aag0851–aag0851.

Nakane, T., Kimanius, D., Lindahl, E., and Scheres, S.H. (2018). Characterisation of molecular motions in cryo-EM single-particle data by multi-body refinement in RELION. Elife 7, 1485.

Nishimura, Y., and Martin, M.A. (2017). Of Mice, Macaques, and Men: Broadly Neutralizing Antibody Immunotherapy for HIV-1. Cell Host Microbe 22, 207–216.

Noe, S.M., Green, M.A., Hogenesch, H., and Hem, S.L. (2010). Mechanism of immunopotentiation by aluminum-containing adjuvants elucidated by the relationship between antigen retention at the inoculation site and the immune response. Vaccine 28, 3588–3594.

Pauthner, M., Havenar-Daughton, C., Sok, D., Nkolola, J.P., Bastidas, R., Boopathy, A.V., Carnathan, D. G., Chandrashekar, A., Cirelli, K.M., Cottrell, C.A., et al. (2017). Elicitation of Robust Tier 2 Neutralizing Antibody Responses in Nonhuman Primates by HIV Envelope Trimer Immunization Using Optimized Approaches. Immunity 46, 1073–1088.e1076.

Petrovas, C., Yamamoto, T., Gerner, M.Y., Boswell, K.L., Wloka, K., Smith, E.C., Ambrozak, D.R., Sandler, N.G., Timmer, K.J., Sun, X., et al. (2012). CD4 T follicular helper cell dynamics during SIV infection. J. Clin. Invest. 122, 3281–3294.

Plotkin, S.A. (2010). Correlates of protection induced by vaccination. Clin. Vaccine Immunol. 17, 1055–1065.

Potter, C.S., Chu, H., Frey, B., Green, C., Kisseberth, N., Madden, T.J., Miller, K.L., Nahrstedt, K., Pulokas, J., Reilein, A., et al. (1999). Leginon: a system for fully automated acquisition of 1000 electron micrographs a day. Ultramicroscopy 77, 153–161.

Price, M.N., Dehal, P.S., and Arkin, A.P. (2010). FastTree 2--approximately maximum-likelihood trees for large alignments. PLoS ONE 5, e9490.

Ramesh, A., Darko, S., Hua, A., Overman, G., Ransier, A., Francica, J.R., Trama, A., Tomaras, G.D., Haynes, B.F., Douek, D.C., et al. (2017). Structure and Diversity of the Rhesus Macaque Immunoglobulin Loci through Multiple De Novo Genome Assemblies. Front Immunol 8, 220–19.

Rerks-Ngarm, S., Pitisuttithum, P., Nitayaphan, S., Kaewkungwal, J., Chiu, J., Paris, R., Premsri, N., Namwat, C., de Souza, M., Adams, E., et al. (2009). Vaccination with ALVAC and AIDSVAX to prevent HIV-1 infection in Thailand. N. Engl. J. Med. 361, 2209–2220.

Richman, D.D., Wrin, T., Little, S.J., and Petropoulos, C.J. (2003). Rapid evolution of the neutralizing antibody response to HIV type 1 infection. Proceedings of the National Academy of Sciences 100, 4144–4149.

Robinson, J.T., Thorvaldsdóttir, H., Winckler, W., Guttman, M., Lander, E.S., Getz, G., and Mesirov, J.P. (2011). Integrative genomics viewer. Nat. Biotechnol. 29, 24–26.

Sanders, R.W., Derking, R., Cupo, A., Julien, J.-P., Yasmeen, A., de Val, N., Kim, H.J., Blattner, C., la Peña, de, A.T., Korzun, J., et al. (2013). A next-generation cleaved, soluble HIV-1 Env trimer, BG505 SOSIP.664 gp140, expresses multiple epitopes for broadly neutralizing but not non-neutralizing antibodies. PLoS Pathog. 9, e1003618.

Sanders, R.W., van Gils, M.J., Derking, R., Sok, D., Ketas, T.J., Burger, J.A., Ozorowski, G., Cupo, A., Simonich, C., Goo, L., et al. (2015). HIV-1 VACCINES. HIV-1 neutralizing antibodies induced by native-like envelope trimers. Science 349, aac4223–aac4223.

Scheres, S.H.W. (2012). RELION: implementation of a Bayesian approach to cryo-EM structure determination. J. Struct. Biol. 180, 519–530.

Schwickert, T.A., Victora, G.D., Fooksman, D.R., Kamphorst, A.O., Mugnier, M.R., Gitlin, A.D., Dustin, M.L., and Nussenzweig, M.C. (2011). A dynamic T cell-limited checkpoint regulates affinity-dependent B cell entry into the germinal center. J. Exp. Med. 208, 1243–1252.

Shi, Y., Hogenesch, H., and Hem, S.L. (2001). Change in the degree of adsorption of proteins by aluminum-containing adjuvants following exposure to interstitial fluid: freshly prepared and aged model vaccines. Vaccine 20, 80–85.

Sorzano, C.O.S., Bilbao-Castro, J.R., Shkolnisky, Y., Alcorlo, M., Melero, R., Caffarena-Fernández, G., Li, M., Xu, G., Marabini, R., and Carazo, J.M. (2010). A clustering approach to multireference alignment of single-particle projections in electron microscopy. J. Struct. Biol. 171, 197–206.

Stewart-Jones, G.B.E., Soto, C., Lemmin, T., Chuang, G.-Y., Druz, A., Kong, R., Thomas, P.V., Wagh, K., Zhou, T., Behrens, A.-J., et al. (2016). Trimeric HIV-1-Env Structures Define Glycan Shields from Clades A, B, and G. Cell 165, 813–826.

Sundling, C., Phad, G., Douagi, I., Navis, M., and Karlsson Hedestam, G.B. (2012). Isolation of antibody V(D)J sequences from single cell sorted rhesus macaque B cells. J. Immunol. Methods 386, 85–93.

Tam, H.H., Melo, M.B., Kang, M., Pelet, J.M., Ruda, V.M., Foley, M.H., Hu, J.K., Kumari, S., Crampton, J., Baldeon, A.D., et al. (2016). Sustained antigen availability during germinal center initiation enhances antibody responses to vaccination. Proc. Natl. Acad. Sci. U.S.a. 113, 201606050–E201606648.

Tas, J.M.J., Mesin, L., Pasqual, G., Targ, S., Jacobsen, J.T., Mano, Y.M., Chen, C.S., Weill, J.-C., Reynaud, C.-A., Browne, E.P., et al. (2016). Visualizing antibody affinity maturation in germinal centers. Science 351, 1048–1054.

Thorvaldsdóttir, H., Robinson, J.T., and Mesirov, J.P. (2013). Integrative Genomics Viewer (IGV): high-performance genomics data visualization and exploration. Brief. Bioinformatics 14, 178–192.

Turner, J.S., Benet, Z.L., and Grigorova, I.L. (2017). Antigen Acquisition Enables Newly Arriving B Cells To Enter Ongoing Immunization-Induced Germinal Centers. J. Immunol. 199, 1301–1307.

Upadhyay, A.A., Kauffman, R.C., Wolabaugh, A.N., Cho, A., Patel, N.B., Reiss, S.M., Havenar-Daughton, C., Dawoud, R.A., Tharp, G.K., Sanz, I., et al. (2018). BALDR: a computational pipeline for paired heavy and light chain immunoglobulin reconstruction in single-cell RNA-seq data. Genome Med 10, 20.

Vander Heiden, J.A., Yaari, G., Uduman, M., Stern, J.N.H., O’Connor, K.C., Hafler, D.A., Vigneault, F., and Kleinstein, S.H. (2014). pRESTO: a toolkit for processing high-throughput sequencing raw reads of lymphocyte receptor repertoires. Bioinformatics 30, 1930–1932.

Victora, G.D., and Wilson, P.C. (2015). Germinal center selection and the antibody response to influenza. Cell 163, 545–548.

Victora, G.D., Schwickert, T.A., Fooksman, D.R., Kamphorst, A.O., Meyer-Hermann, M., Dustin, M.L., and Nussenzweig, M.C. (2010). Germinal center dynamics revealed by multiphoton microscopy with a photoactivatable fluorescent reporter. Cell 143, 592–605.

Vigdorovich, V., Oliver, B.G., Carbonetti, S., Dambrauskas, N., Lange, M.D., Yacoob, C., Leahy, W., Callahan, J., Stamatatos, L., and Sather, D.N. (2016). Repertoire comparison of the B-cell receptor-encoding loci in humans and rhesus macaques by next-generation sequencing. Clin Transl Immunology 5, e93.

Voss, N.R., Yoshioka, C.K., Radermacher, M., Potter, C.S., and Carragher, B. (2009). DoG Picker and TiltPicker: software tools to facilitate particle selection in single particle electron microscopy. J. Struct. Biol. 166, 205–213.

Watson, C.T., and Breden, F. (2012). The immunoglobulin heavy chain locus: genetic variation, missing data, and implications for human disease. Genes Immun. 13, 363–373.

Watson, C.T., Glanville, J., and Marasco, W.A. (2017). The Individual and Population Genetics of Antibody Immunity. Trends Immunol. 38, 459–470.

Wei, X., Decker, J.M., Wang, S., Hui, H., Kappes, J.C., Wu, X., Salazar-Gonzalez, J.F., Salazar, M.G., Kilby, J.M., Saag, M.S., et al. (2003). Antibody neutralization and escape by HIV-1. Nature 422, 307–312.

Weissburg, R.P., Berman, P.W., Cleland, J.L., Eastman, D., Farina, F., Frie, S., Lim, A., Mordenti, J., Peterson, M.R., Yim, K., et al. (1995). Characterization of the MN gp120 HIV-1 Vaccine: Antigen Binding to Alum. Pharmaceutical Research 12, 1439–1446.

West, A.P., Scharf, L., Scheid, J.F., Klein, F., Bjorkman, P.J., and Nussenzweig, M.C. (2014). Structural insights on the role of antibodies in HIV-1 vaccine and therapy. Cell 156, 633–648.

Yamamoto, T., Lynch, R.M., Gautam, R., Matus-Nicodemos, R., Schmidt, S.D., Boswell, K.L., Darko, S., Wong, P., Sheng, Z., Petrovas, C., et al. (2015). Quality and quantity of TFH cells are critical for broad antibody development in SHIVAD8 infection. Sci Transl Med 7, 298ra120–298ra120.

Ye, J., Ma, N., Madden, T.L., and Ostell, J.M. (2013). IgBLAST: an immunoglobulin variable domain sequence analysis tool. Nucleic Acids Res. 41, W34–W40.

Yeh, C.-H., Nojima, T., Kuraoka, M., and Kelsoe, G. (2018). Germinal center entry not selection of B cells is controlled by peptide-MHCII complex density. Nat Commun 9, 928.

Zhou, T., Doria-Rose, N.A., Cheng, C., Stewart-Jones, G.B.E., Chuang, G.-Y., Chambers, M., Druz, A., Geng, H., McKee, K., Kwon, Y.D., et al. (2017). Quantification of the Impact of the HIV-1-Glycan Shield on Antibody Elicitation. Cell Rep 19, 719–732.

